# Age-specific and compartment-dependent changes in mitochondrial homeostasis and cytoplasmic viscosity in mouse peripheral neurons

**DOI:** 10.1101/2023.09.18.558344

**Authors:** James N. Sleigh, Francesca Mattedi, Sandy Richter, Emily Annuario, Kristal Ng, I. Emilie Steinmark, Iveta Ivanova, István L. Darabán, Parth P. Joshi, Elena R. Rhymes, Shirwa Awale, Gokhan Yahioglu, Jacqueline C. Mitchell, Klaus Suhling, Giampietro Schiavo, Alessio Vagnoni

## Abstract

Mitochondria are dynamic bioenergetic hubs that become compromised with age. In neurons, declining mitochondrial axonal transport has been associated with reduced cellular health. However, it is still unclear to what extent the decline of mitochondrial transport and function observed during ageing are coupled, and if somal and axonal mitochondria display compartment-specific features that make them more susceptible to the ageing process. It is also not known whether the biophysical state of the cytoplasm, thought to affect many cellular functions, changes with age to impact mitochondrial trafficking and homeostasis. Focusing on the mouse peripheral nervous system, we show that age-dependent decline in mitochondrial trafficking is accompanied by reduction of mitochondrial membrane potential and intra-mitochondrial viscosity, but not calcium buffering, in both somal and axonal mitochondria. Intriguingly, we observe a specific increase in cytoplasmic viscosity in the neuronal cell body, where mitochondria are most polarised, which correlates with decreased cytoplasmic diffusiveness. Increasing cytoplasmic crowding in the somatic compartment of DRG neurons grown in microfluidic chambers reduces mitochondrial axonal trafficking, suggesting a mechanistic link between the regulation of cytoplasmic viscosity and mitochondrial dynamics. Our work provides a reference for studying the relationship between neuronal mitochondrial homeostasis and the viscoelasticity of the cytoplasm in a compartment-dependent manner during ageing.

## INTRODUCTION

Ensuring precise distribution and functionality of mitochondria is essential for maintaining cellular homeostasis. The complex architecture and long distances between nerve terminals and soma of neurons cause these cells to be highly dependent on mechanisms for long-range cytoskeletal transport of mitochondria and on continuous mitochondrial supply and functionality along their extended cellular architecture. This is a particularly critical challenge for neurons, which are post-mitotic and can live throughout the adult life of an organism. It is therefore not surprising that mitochondrial dysfunction is a recognised hallmark of ageing and neurodegeneration (López-Otín et al., 2023).

We have previously shown that axonal transport of mitochondria decreases in the peripheral nervous system of ageing *Drosophila melanogaster* (Vagnoni et al., 2016) and that neuronal ageing phenotypes, such as oxidative stress and focal protein accumulation, can be manipulated by modulation of mitochondrial transport (Vagnoni & Bullock, 2018; Vagnoni et al., 2016). Mitochondrial motility was shown to be downregulated during ageing to varying degrees also in *C. elegans* motor neurons and in the mouse central nervous system neurons *in vivo* (Mattedi & Vagnoni, 2019; Morsci et al., 2016; Takihara et al., 2015). Despite the consistent observation that mitochondrial trafficking is slowed down during ageing, it remains unclear whether this directly correlates with functional changes of the mitochondria.

The biophysical properties of the neuronal cytoplasm are under increasing scrutiny in the context of neurodegeneration (Alberti & Hyman, 2016; Elbaum-Garfinkle & Brangwynne, 2015), although the effect of age on these properties is less well studied. Whether fundamental properties of the neuroplasm, such as viscosity and diffusion, undergo significant changes that can affect intracellular cargo trafficking and functionality during ageing is currently unknown.

A study that combines analyses of the motile behaviour of neuronal cargoes and their functional state with the evaluation of the biophysical properties of the cytoplasm in young and old neurons has been missing, partly due to lack of suitable experimental systems to perform this type of work and the challenge of studying these processes in live neurons throughout ageing. Interrogating cellular processes in animal models *in vivo* is indispensable for ageing studies, although gaining mechanistic insight can be difficult in this context (Sleigh et al., 2017a). For example, neuronal cells *in situ* often have their soma, axons and dendrites embedded in different tissues and organs and so studying functional compartmentalisation *in vivo* becomes challenging. In an ageing context, being able to complement *in vivo* studies with a species-specific *in vitro* model of neuronal ageing would be particularly advantageous.

In this study, by focusing on the mouse peripheral nervous system, we filled this gap through comparative analyses of mitochondrial trafficking in dorsal root ganglion (DRG) neurons *in vitro* and sciatic nerve axons *in vivo* from both young and aged mice. Using the DRG model, we also monitored other important aspects of mitochondrial biology within neurons, including mitochondrial membrane potential, calcium uptake and intra-mitochondrial viscosity, and we revealed age- and compartment-dependent alterations in these properties. We expanded on these data by quantifying intracellular viscosity and diffusion in the neuronal cytoplasm of the somal and axonal compartments and uncover a mechanistic link between increased crowding and viscosity in the cell bodies and reduced mitochondrial axonal trafficking. With this work, we propose a novel model that correlates mitochondrial homeostasis with the viscoelasticity of the neuronal cytoplasm during ageing.

## RESULTS

### Mitochondrial trafficking declines with age in mouse peripheral neurons both *in vitro* and *in vivo*

To better characterise the effect of ageing on mitochondrial trafficking in mouse peripheral neurons, we cultured sensory neurons from DRG of young (3-4 months) and old (19-22 months) mice (Fig. 1A) and analysed mitochondrial axonal transport by live imaging with MitoTracker. DRG neurons grow robustly in culture after dissection, retain their age-dependent transcriptional profile (Gumy et al., 2011) and have previously been used to study axonal cargo trafficking during ageing (Mellone et al., 2013; Stavoe et al., 2019). Time-lapse recording and quantification of mitochondrial axonal transport showed abundant mitochondrial movement in both the anterograde and retrograde directions, with a small proportion of mitochondria displaying bidirectional motility (Fig. 1B-C, Fig. S1A-B) (Klinman & Holzbaur, 2015; Lin et al., 2017; Mellone et al., 2013; Pandey et al., 2022). Neurons obtained from old mice showed a strong reduction in the percentage of moving mitochondria (Fig. 1B-C, Fig. S1A-B, Movie 1), but no significant change in the number of mitochondria present in axons (Fig. 1D), which correlated with an increase in their overall size (Fig. 1E-G, Fig. S1C-D). Moreover, the speed and displacement of these organelles were largely unaffected by age, both in the anterograde (Fig. 1H) and retrograde directions (Fig. 1I).

**Figure 1.**
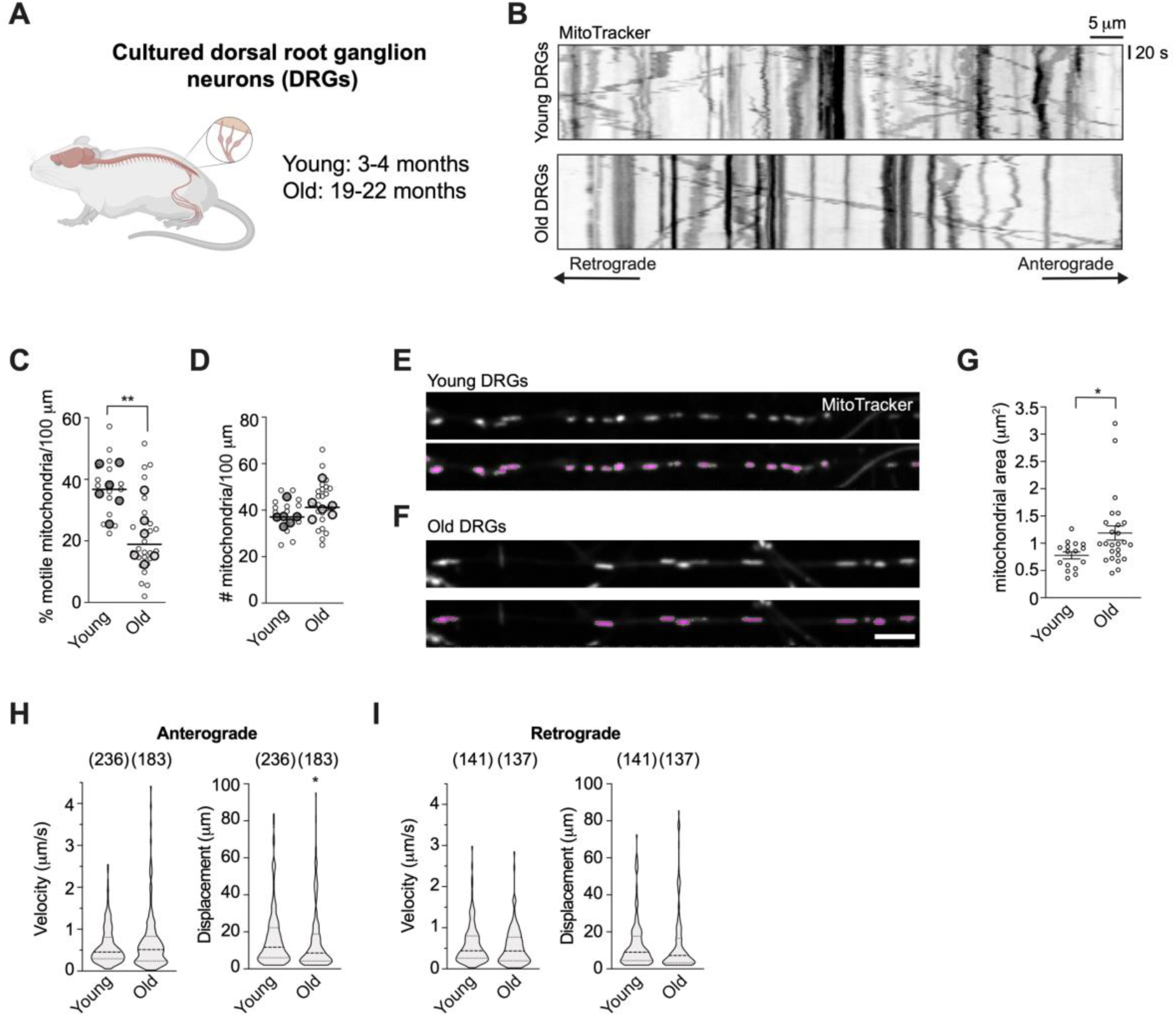
Mitochondrial transport is reduced in aged DRG neurons in culture. **(A)** DRGs were dissected from the spinal cord of 6 young (P72-107, 3 females and 3 males) and 6 old (P560-637, 3 females and 3 males) mice and dissociated neurons were cultured *in vitro.* **(B)** Representative kymographs of mitochondria moving in the axons of DRG neurons dissected from young and old animals. **(C-D)** The proportion of motile mitochondria (C), but not the mitochondrial mass (D), is strongly reduced during ageing in the DRG neuron axons. Individual data points representing the miceused (bigger circles with line depicting the median) are overlayed on data points (smaller circles) representing individual fields of view (FOV) imaged (1-3 axons/FOV). **(E-F)** Representative frames of MitoTracker-stained mitochondria in the axons of cells extracted from young (E) and old animals (F). Bottom panels show the thresholded mitochondrial area (magenta) used for quantitative analysis of morphology. Scale bar: 5 μm. **(G)** Quantification of mitochondrial area in young and old cells. Individual data points represent the fields of view (FOV) imaged (1-3 axons/FOV) and are shown as mean ± SEM. **(H-I)** Violin plot distributions show that anterograde (H) and retrograde (I) mitochondrial velocity and displacement remain largely stable during ageing, albeit with a subtle but significant reduction in anterograde displacement in aged neurons. In brackets, number of mitochondria. Unpaired student’s t-test (C-D), Mann-Whitney test (G-I). * p < 0.05; ** p < 0.01.

To assess whether the age-dependent decline in mitochondrial transport observed in cultured DRGs was preserved in peripheral neurons *in vivo*, we analysed mitochondrial axonal transport in intact sciatic nerve axons of anaesthetised MitoMice (Fig. 2A), in which neuronal mitochondria are marked by cyan fluorescent protein (CFP) (Fig. 2B-C and Movie 2) (Misgeld et al., 2007). Old mice displayed a clear reduction in mitochondrial motility (Fig. 2D and Movie 2), which was accompanied by a reduction in the overall number of axonal mitochondria (Fig. 2E) and a stark increase in mitochondrial area (Fig. 2F, Fig. S2A-B). When the proportion of mitochondrial motility was assessed, no difference between young and old neurons was identified (Fig. S2C) in either anterograde or retrograde directions (Fig. S2D). This indicates that the decline in the number of motile mitochondria might not be due to a net change in the proportion of moving mitochondria, rather a potential imbalance in mitochondrial biogenesis and/or turnover that leads to altered mitochondrial content in axons. While the motile properties of the retrograde-moving mitochondria remained unchanged with age (Fig. 2H), the speeds and displacement of the fewer anterograde-moving mitochondria were surprisingly upregulated in older mice (Fig. 2G), contrasting with what observed in cultured DRG neurons. Interestingly, the distributions of mitochondrial speed and displacement measured in the sciatic nerve and cultured neurons were significantly different (Fig. 1H-I, 2G-H, and Table S1), potentially indicating a different regulation of the molecular motor machinery responsible for the movement of these organelles *in vivo* and *in vitro*. Overall, these data show that mitochondrial axonal trafficking is affected by age in peripheral nerves both *in vitro* and *in vivo*, although the underlying mechanisms may differ.

**Figure 2.**
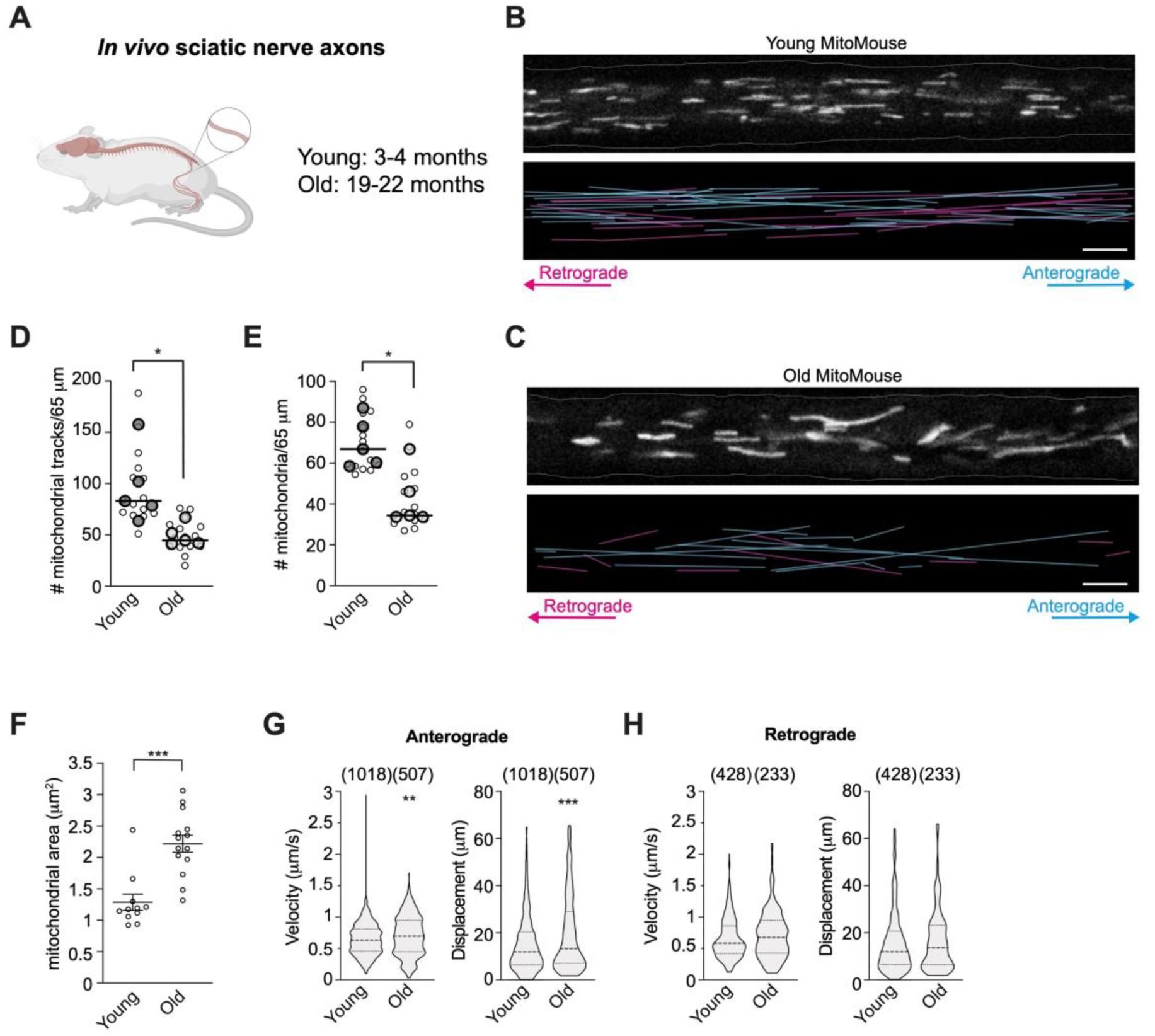
Mitochondrial transport decreases with age in the sciatic nerve *in vivo*. 5 young (P72-107, 3 females and 2 males) and 5 old (P560-637, 2 females and 3 males) MitoMice were anaesthetised and the sciatic nerve exposed for time-lapse imaging of mitochondrial axonal transport. **(B-C)** Still frames from movies showing CFP-marked mitochondria (grey, top panels) in single sciatic nerve axons from young and old (C) MitoMice. The bottom panels depict the traces of mitochondria moving in the anterograde and retrograde directions (cyan and magenta, respectively) in the corresponding movies. Scale bars: 5 μm. **(D-E)** Both the number of motile mitochondria and the total number of mitochondria present in the axons of the sciatic nerve are strongly reduced during ageing. Individual data points representing the mice used (bigger circles with line depicting median) are overlayed on data points (smaller circles) representing individual sciatic nerve axons. **(F)** Quantification of mitochondrial area in young and old sciatic nerves. Individual data points represent the sciatic nerve axons and are shown as mean ± SEM. **(G-H)** Violin plot distributions show that mitochondrial anterograde velocity and displacement are both upregulated in old neurons (G), while the motile properties of the retrograde-moving mitochondria are not affected by age (H). In brackets, number of mitochondria. Mann-Whitney test (D-H). * p < 0.05; ** p < 0.01; *** p < 0.001.

### Mitochondrial membrane potential is compartment-dependent and decreases with age and declining mitochondrial viscosity

To understand if the trafficking defects are associated with altered mitochondrial functionality, we first asked whether the membrane potential (ΔΨ_m_) of these organelles was altered between young and old DRG neurons. We stained the neurons with a low concentration of the live dye tetramethylrhodamine methyl ester (TMRM), which accumulates in active mitochondria, as well as MitoTracker Green (MTG) to label all mitochondria (Fig. 3A-D). Calculating the ratio of TMRM to MTG fluorescence has previously been used to determine ΔΨ_m_ in live cells, where a higher ratio indicates more polarised mitochondria (Mitra & Lippincott-Schwartz, 2010). Using this method here reveals that the membrane potential of neuronal mitochondria decreases with age (Fig. 3E and Fig. S3A-B) and that the axonal compartment is characterised by mitochondria with substantially lower membrane potential, independently of age (Fig. 3F-G). Using the ratiometric dye JC-1, we confirmed that axonal mitochondria are more depolarised than somal mitochondria (Fig. S3C-D), further validating our protocol to estimate ΔΨ_m_, using the TMRM/MTG ratio.

**Figure 3.**
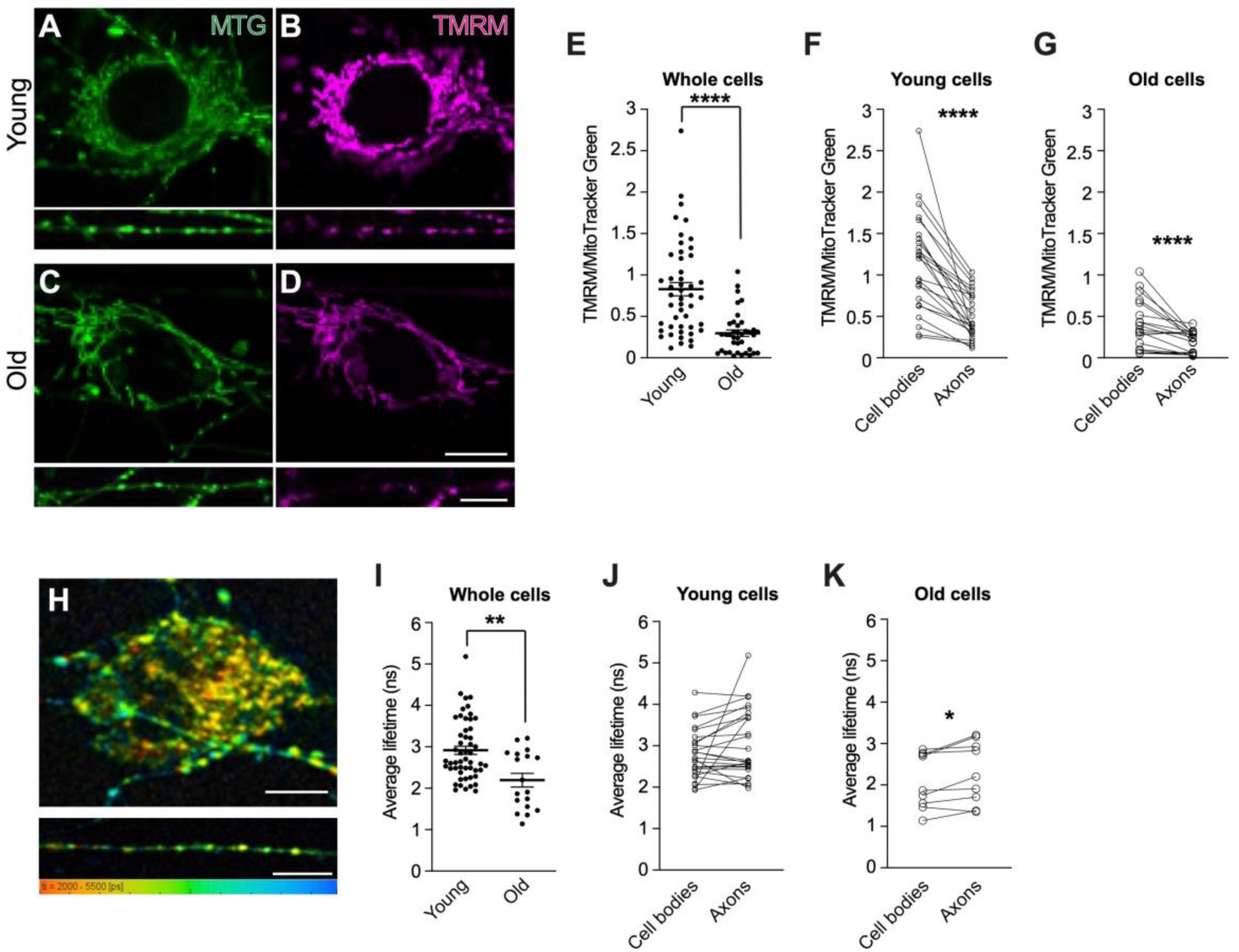
Mitochondrial membrane potential and viscosity decline during ageing. **(A-D)** Representative images of young (A-B) and aged (C-D) DRG neuron cell bodies (top panels) and axons (bottom panels), stained with TMRM and MitoTracker Green (MTG). Scale bars: 10 μm. **(E-G)** Quantification of the ratio between TMRM and MTG in neuronal cell bodies and axons combined indicates that the ΔΨ_m_ declines during ageing at cellular level (E) with axonal mitochondria displaying consistently reduced ΔΨ_m_ in both young (F) and old (G) cells. **(H)** Representative images of a DRG neuron cell body (top panel) and axon (bottom panel) stained with FMR-1 to capture mitochondrial viscosity. Rainbow colour scale indicates fluorescence lifetime range (2,000-5,500 ps) in which orange/red and cyan/blue hues correspond to areas of lower and higher viscosity, respectively. Scale bars: 15 μm. **(I)** Overall mitochondrial viscosity is decreased in old cells, as indicated by quantification in neuronal cell bodies and axons combined. **(J-K)** There is no significant difference between somal and axonal mitochondrial viscosity in young DRG neurons (J), while mitochondrial viscosity is slightly higher in old DRG axons (K). In E-G and I-K, data points represent the cell bodies and axons imaged. Each neuron analysed provides both cell body and axons to the analysis. DRGs were dissected from 3 young (P49-67, 1 female and 2 males) and 3 old (P672-679, 2 females and 1 male) mice for TMRM experiments (A-G), and from 3 young (P81-84, 2 females and 1 male) and 2 old (P715, 1 female and 1 male) mice for FLIM experiments (H-K). Data are shown as mean ± SEM. Mann-Whitney test (E, I), paired student’s t-test (F, K), Wilcoxon test (G, J). * p < 0.05; ** p < 0.01; **** p < 0.0001.

It is possible that decreased trafficking and altered membrane potential of mitochondria during ageing are mechanistically linked, although the two phenotypes could also manifest independently. To test if decreased mitochondrial trafficking has a direct effect on membrane potential in DRGs, we used an RNAi construct to reduce the abundance of Trak1, which is essential to drive mitochondrial motility in axons (van Spronsen et al., 2013). Trak1 knockdown in young DRG neurons reduced mitochondrial transport (Fig. S3E) but had no impact on membrane potential (Fig. S3F-G). This indicates that acute reduction of mitochondrial axonal transport does not directly affect ΔΨ_m_, suggesting that the two phenotypes in the aged neurons are not mechanistically linked. Nevertheless, we cannot exclude that chronic reduction of mitochondrial transport during ageing or a time-specific reduction in older neurons could have a knock-on effect on the maintenance of ΔΨ_m_.

Manipulations that decrease mitochondrial membrane potential have been shown to affect the viscosity of these organelles in HeLa and COS7 cells (Chambers et al., 2018; Chen et al., 2020; Guo et al., 2022; Steinmark et al., 2019), while altered mitochondrial membrane viscosity has been observed in brains of people affected by Alzheimer’s disease (Mecocci et al., 1997) and in murine neurons during neurodegeneration and ageing (Eckmann et al., 2014; Kuter et al., 2016). We therefore asked if the age-dependent changes in ΔΨ_m_ that we observe in DRGs might be accompanied by changes in the viscosity of the mitochondria. To do so, we performed fluorescence lifetime imaging microscopy (FLIM) of neurons incubated with the BODIPY-based fluorescent molecular rotor FMR-1 targeted to the mitochondrial matrix (Fig. 3H) (Steinmark et al., 2019). Fluorescence lifetime of FMR-1 responds to changes in viscosity environments, where a longer fluorescence lifetime indicates higher viscosity. Neuronal mitochondria display a heterogeneous viscosity profile both in the cell bodies and axons (Figure 3H), in accordance with previous work carried out in HeLa and COS7 cells (Chambers et al., 2018; Steinmark et al., 2019). Similar to ΔΨ_m_, average mitochondrial viscosity decreases during ageing as indicated by decreased lifetime (Fig. 3I and Fig. S3H-I). In this case, however, the viscosity of the mitochondria remains largely unchanged between the cell bodies and axons in both young and old neurons (Fig. 3J-K). These results suggest that while the striking difference in ΔΨ_m_ between somal and axonal mitochondria is unlikely to be coupled to organelle viscosity (as this remains constant across compartments), the decline of ΔΨ_m_ and mitochondrial viscosity during age may be linked.

### Ca^2+^ buffering is unaffected in DRG neurons cultured from young and old mice

Loss of ΔΨ_m_ has long been thought to negatively affect Ca^2+^ accumulation in mitochondria (Brini et al., 1999; Duchen, 2000) and, in mouse sensory neurons, [Ca^2+^]_m_ uptake causes a transient decrease in ΔΨ_m_ (Duchen, 2000). Increased mitochondrial viscosity was also suggested to correlate with augmented cytosolic calcium in HeLa cells (Steinmark et al., 2019). We therefore asked whether the age-dependent changes in the ΔΨ_m_ and mitochondrial viscosity are accompanied by alterations in [Ca^2+^]_m_ uptake. We depolarised DRG cultures by transient stimulation with 50 mM KCl and measured the fluorescent signal of mitochondrial and cytosolic Ca^2+^ using the Rhod-2-AM and Fluo4-AM dyes, respectively (De vos et al., 2012; Duchen, 2000) (Fig. S4A-J). Combined measurements from both the somal and axonal compartments showed that there was no overall difference in the timing and abundance of mitochondrial Ca^2+^ uptake and release (Rhod-2-AM) between young and old cells (Fig. S4K,M,O). Likewise, we did not detect any significant difference in the dynamics of cytoplasmic Ca^2+^ (Fluo4-AM) between young and old neurons following neuronal stimulation (Fig. S4L,N,P). Taken together, these data show that age does not affect Ca^2+^ homeostasis in cultured DRGs.

### Age-dependent increase in cytoplasmic viscosity is inversely correlated with diffusiveness in neuronal cell bodies but not in axons

Previous studies have shown that hypertonic treatments, which acutely increase molecular crowding and cytoplasmic viscosity, affect cytoskeletal dynamics in fission yeast (Molines et al., 2022), alter ER-to-Golgi transport in COS7 cells (Lee & Linstedt, 1999) and abolish endosomal motility in mammalian immune cells (Nunes et al., 2015). To test whether decreased mitochondrial transport in ageing DRG neurons correlates with a change in fluid-phase viscosity of the cytoplasm, we performed time-resolved fluorescence <u>a</u>nisotropy <u>im</u>aging (TR-FAIM, Fig. 4A) (Clayton et al., 2002; Siegel et al., 2003). TR-FAIM measures the depolarisation of fluorescence emission of a fluorescein derivative (2′,7′-bis-(2-carboxyethyl)-carboxyfluorescein, BCECF) caused by rotational Brownian motion, and so it is used to obtain information about the viscosity of the cellular microenvironment (Dix & Verkman, 1990). Anisotropy decay measurements and fitting (Fig. S5A and see Methods) showed that the rotational correlation time of the BCECF is significantly reduced in older cells leading to slower depolarisation (Fig. 4B), which is indicative of higher cytoplasmic viscosity in old DRGs (Fig. 4C). Because of the specialised architecture of neurons, we hypothesised that the somal and axonal compartments might comprise different viscosity environments. Measurement of the anisotropy decay in cell bodies and axons revealed that, although the two compartments have similar viscosity values in young cells (Fig. 4D, F and Fig. S5B), the viscosity of the cell bodies undergoes a significant increase in aged cells (Fig. 4E-F and Fig. S5C-D), whereas the axonal viscosity remains unchanged during age (Fig. 4D-F and Fig. S5B-C, E). These results suggest that axonal viscosity alone is unlikely to explain the observed decrease in mitochondrial axonal transport.

**Figure 4.**
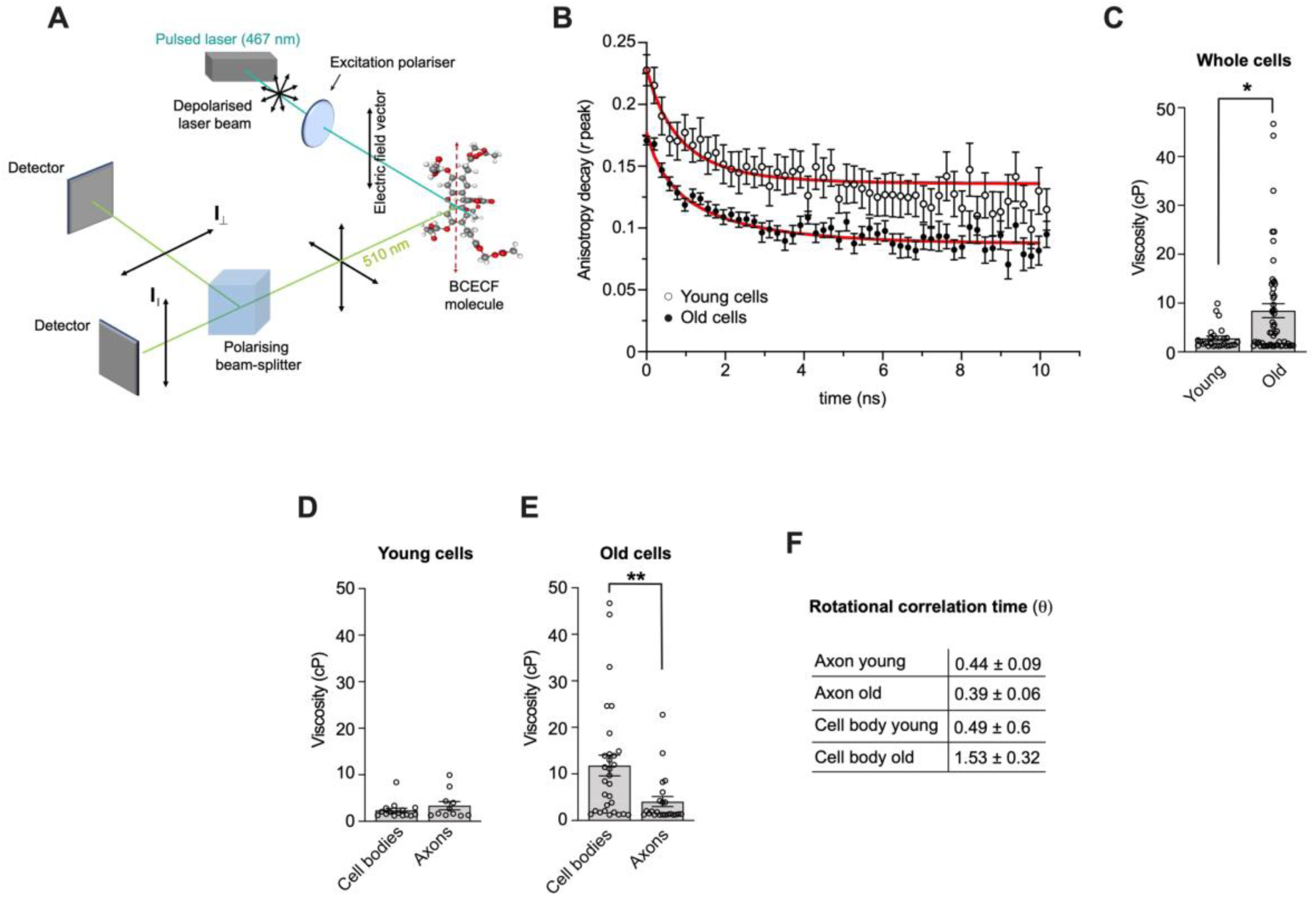
The cytoplasmic viscosity of aged DRG neurons in culture increases in cell bodies but not in axons. **(A)** Schematic of the fluorescence anisotropy optical set up. A 467 nm depolarised pulsed laser beam (cyan) is linearly polarised before exciting BCECF molecules with a transition dipole (dashed red line) parallel to the electric field vector of the light. The 510 nm emitted light (green) is passed into a polarising beam splitter to two detectors for light with either parallel (I∥) or perpendicular (I⊥) emission polarisation. The rotational correlation time (θ) of the BCECF molecule is defined as the time taken to rotate one radian (during the fluorescence lifetime). **(B)** Fluorescence anisotropy decay and fit (see Methods, equation 3) of BCECF fluorescence obtained from perpendicular and parallel decays (according to equation 1, see Methods) from young and old DRG neurons. Older cells display a slower decay, indicative of higher cytoplasmic viscosity. Red lines show the exponential decay fit and circles the individual time points. **(C)** Viscosity values obtained from anisotropy decay analysis show increased viscosity in old DRG neurons. Circles show individual cellular compartments. Relative to (B). **(D-E)** The somal and axonal compartments display similar viscosity values in young DRG neurons (D, p = 0.5065), whereas the cytoplasmic viscosity of old DRG neurons is significantly higher in the somal compartment. Circles depict individual cellular compartments. **(F)** Rotational correlation time (θ) values relative to somal and axonal anisotropy data. Note the average θ is proportional to viscosity. Data are shown as mean ± SEM, Mann-Whitney test. * p < 0.05; ** p < 0.01. DRGs for the anisotropy experiments were obtained from 2 young (P80) and 3 old (P709-712) male mice.

Fluid-phase viscosity affects the diffusiveness of the cytoplasmic milieu, as determined by the Stokes-Einstein law (Miller & Walker, 1924). To gain more insight into cytoplasmic diffusiveness in our *in vitro* cell model of ageing, we performed rheological studies by measuring the diffusion rates of genetically-encoded multimeric nanoparticles (GEMs) (Delarue et al., 2018). GEMs can be visualised as bright spherical particles of uniform shape and size that are suitable for microrheological studies in different cellular contexts (Delarue et al., 2018; Molines et al., 2022; Shu et al., 2022). We transduced DRGs with a lentiviral vector encoding sapphire-tagged GEMs of 40 nm diameter (Fig. 5A, D) and tracked particle movement in cell bodies and axons (Fig. 5B-C, E-F and Movie 3-4). Single particle tracking and MSD analysis showed that GEM movement is consistent with sub-diffusive motion (Fig. 5G and Fig. S6A), in accordance with previous reports in non-neuronal cell types (Delarue et al., 2018; Molines et al., 2022). In both young and old neurons, the diffusion rate of axonal GEMs is significantly lower than the rate of GEMs freely diffusing in cell bodies (Fig. 5G-I and Fig. S6), suggesting that the specialised axonal architecture poses a significant physical constraint for particle diffusion. The diffusion of somal GEMs is reduced in aged cultures (Fig. 5G-I), in accordance with the observed increase in viscosity. Interestingly, although the axonal viscosity does not change during ageing (Fig. 4D-F, Fig. S5B-C, E), GEMs display a small, albeit significant, reduction in diffusiveness in aged culture (Fig. 5G-I), indicating that mechanisms other than viscosity regulate axonal diffusion during ageing.

**Figure 5.**
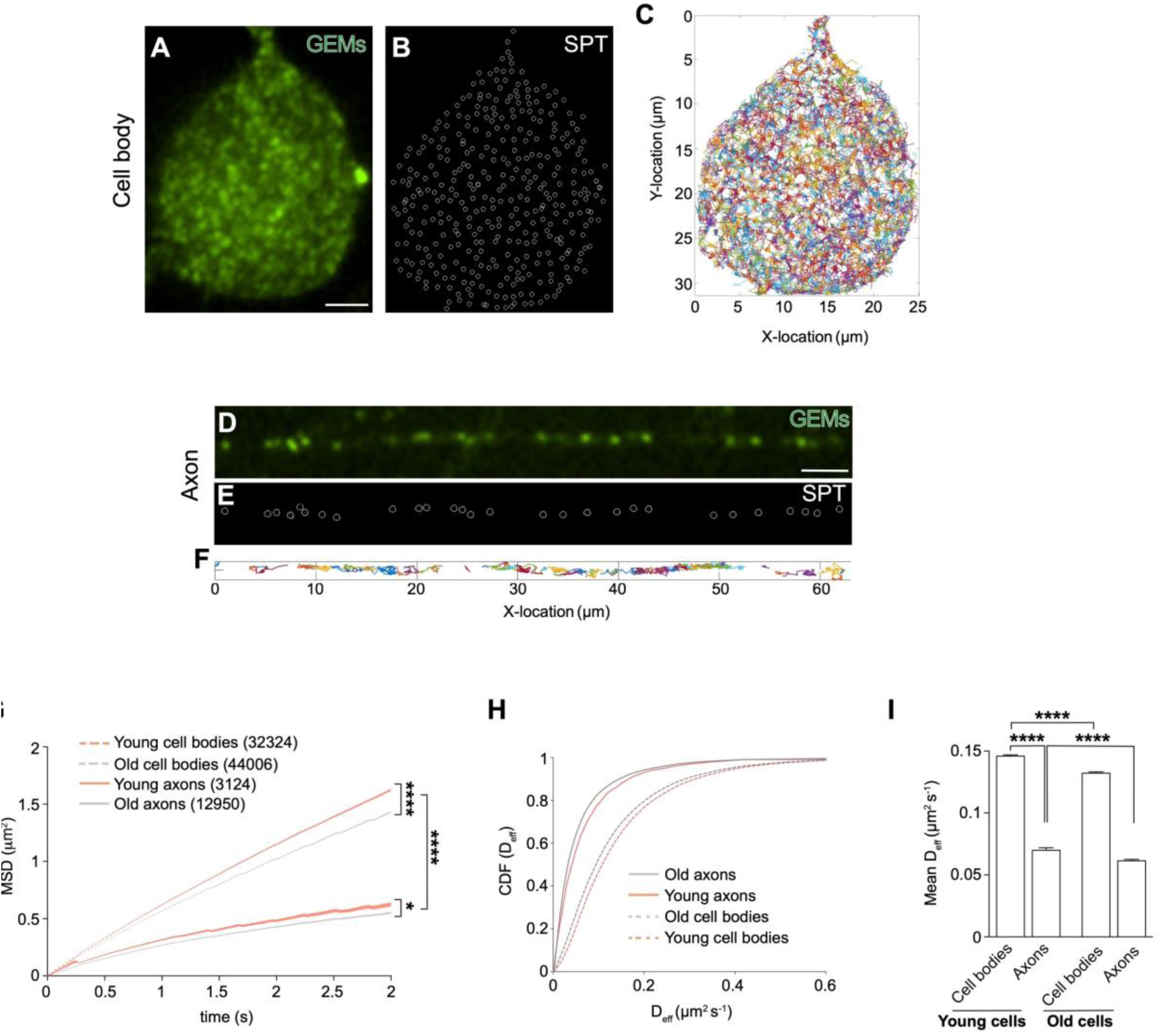
The diffusiveness of the neuronal cytoplasm decreases in aged cultured neurons. **(A-F)** Representative images of a cell body (A-C) and axon (D-F) of a DRG neuron transduced with lentiviral vectors expressing GEMs. In (B) and (E), single particle tracking (SPT) markers are overlayed onto the cell body and axons, respectively. (C) and (F) show the colour-coded GEMs trajectories tracked during 30 s of imaging of and (E), respectively. Scale bars: 5 μm. **(G)** Mean square displacement (MSD) of GEM tracks in cell bodies and axons of young and old cells. Data are plotted as mean ± SEM, Kolmogorov-Smirnov test. The motion of GEMs is captured by a sub-diffusion model described by the equation *MSD (t) = 4Dt^α^* (young cell bodies: R = 0.999, old cell bodies: R = 1, young axons: R = 0.997, old axons: R = 1). In brackets, number of tracks analysed from 61 cell bodies and 170 axons. **(H)** Cumulative distribution function (CDF) of GEMs diffusion coefficient (D_eff_) in young and old DRG axons and cell bodies, relative to (G). **(I)** Mean diffusion coefficient in the cell body and axonal compartments of young and old cells. Relative to (H). DRGs for the microrheology experiments were obtained from 2 young (P49-70) male mice and 2 old (P726-757, 1 female and 1 male) mice. Data are presented as mean ± SEM, Kruskal-Wallis with Dunn’s multiple comparisons test. * p < 0.05; **** p < 0.0001.

Based on these experiments, we conclude that the behaviour of cytoplasmic fluid-phase viscosity during ageing is defined by the cellular compartment, with potentially distinct effects on somal and axonal biology. Indeed, while increased viscosity likely contributes to reduced cytoplasmic diffusion in the neuronal cell bodies, axonal viscosity is not a driver of the decreased diffusiveness of the axonal cytoplasm nor of the reduced active mitochondrial axonal transport during ageing.

### Challenging DRG neuron cell bodies with crowding agents reduces axonal mitochondrial transport

It is conceivable that the stark increase in cytoplasmic viscosity observed in the soma of aged DRGs is sufficient to impact mitochondrial dynamics, including mitochondrial export into the axon, reducing axonal transport rates in old cells. To test this hypothesis, we set out to manipulate the viscosity specifically within somal compartments of young DRG neurons cultured in microfluidic chambers (MFCs)(Fig. 6A, Fig. S7A) to mimic the increased viscosity observed in the aged DRGs. We took advantage of the physical compartmentalisation afforded by MFCs and devised a pulse-chase experiment whereby the somal compartment was stained for 30 min with MitoTracker Green (MTG) following incubation with low doses of either polyethylene glycol (PEG) or sorbitol (Fig. 6B), two crowding agents routinely used to increase intracellular viscosity (Chambers et al., 2018; Molines et al., 2022). The axonal compartment was visualised by staining with CellMask Orange (CMask) to assess axonal growth (Fig. 6B-E). After dyes wash-off, we immediately imaged mitochondrial axonal transport in the chamber microgrooves. We reasoned that any MTG-positive mitochondria observed in the axons would have originated from the somal compartment after being stained during the 30-min pulse. Low-magnification images consistently showed that while MTG-positive mitochondria can travel long distances into the axons-filled microgrooves of untreated cells to reach the synaptic compartments, PEG and sorbitol-treated cells only display faint MTG stain in the proximal axons (Fig. 6C-E). High-resolution time-lapse imaging and quantification of mitochondrial axonal transport reveals a drastic reduction of mitochondrial export from the somal into the axonal compartment in both PEG and sorbitol-treated cells, compared to controls, as indicated by transport fluxes in the initial segment of the microgrooves (Fig. 6F-G, Movie 5). As a result, mitochondrial distribution and occupancy of the axonal space is strongly reduced by these treatments (Fig. S7B, Movie 5). Interestingly, the average velocity of mitochondria is also significantly reduced after sorbitol treatment (Fig. 6H). This suggests the higher viscosity environment of the cell soma induces a lasting effect on the molecular components driving mitochondrial trafficking, which cannot be reversed by translocating into the less crowded axonal environment.

**Figure 6.**
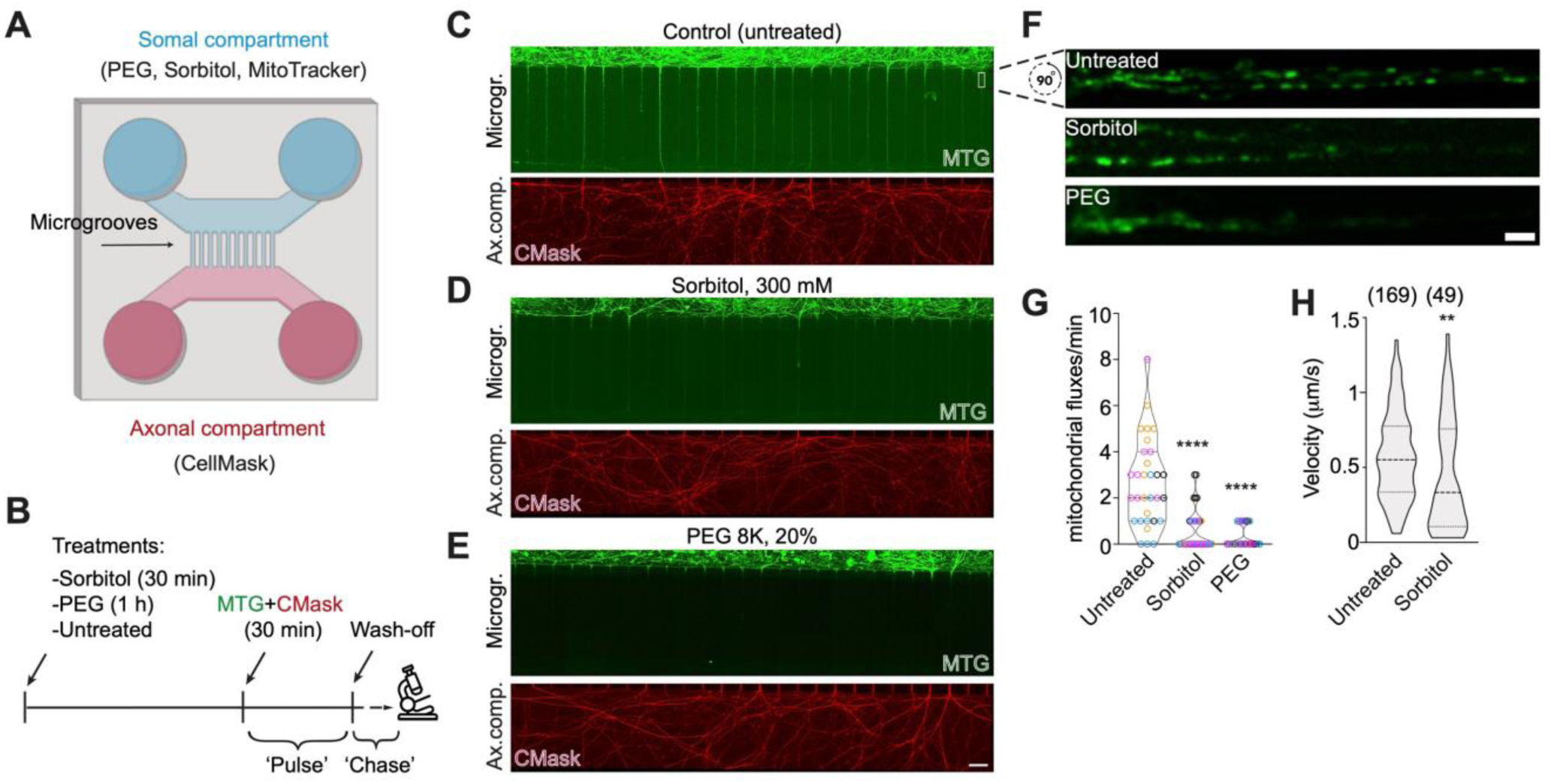
Increasing crowding and viscosity in DRG cell bodies is sufficient to reduce mitochondrial axonal transport. **(A)** Schematic depicting the microfluidic chamber (MFC) used and the treatments specifically applied to the somal and axonal compartments. **(B)** Timeline of the experimental setup, starting with sorbitol or PEG treatment, followed by a ‘pulse’ consisting of staining with live dyes, and a ‘chase’ (i.e. microscopy observation). **(C-E)** Representative images of MFCs in which the somal compartment was either left untreated (C) or treated with sorbitol (D) or PEG (E). Top panels: MFC microgrooves (Microgr.) show the extent of transport from the soma of MitoTracker (MTG)-stained mitochondria during the ‘pulse’ period. Bottom panels: correspondent axonal compartment (Ax. comp.) stained with CellMask (CMask). Scale bar: 100 μm. **(F)** Representative higher magnification of an area of the microgrooves typically used for quantification of mitochondrial motility, indicated by a white rectangle in (C). Scale bar: 5 μm. **(G)** Quantification of mitochondrial transport fluxes (see Methods) shows strong decline in sorbitol and PEG-treated DRGs. Datapoints in the violin plots represent individual microgrooves with each colour indicating a different animal (4 females, P45-86), Kruskal-Wallis with Dunn’s multiple comparisons test. **(H)** Combined axonal mitochondrial velocities are reduced in sorbitol-treated neurons. Due to the very low number of mitochondrial runs in PEG-treated axons, this sample has not been used for statistical analysis. ** p < 0.01; **** p < 0.0001.

The DRG cell body area was not noticeably affected by the treatments with these crowding agents (Fig. S7C-D), at least at the concentrations and incubation times used in this study. This is in line with the unchanged cell body area observed in young and old DRGs (Fig. S6D) and indicates our relatively mild treatments do not trigger an overt osmotic stress. In conclusion, we demonstrate that viscosity manipulations restricted to the neuronal cell bodies, mimicking an aged state, are sufficient to affect mitochondrial transport rates in the axonal compartment.

## DISCUSSION

In this study, we report several new findings that define mitochondrial homeostasis and cytoplasmic biophysics in aging neurons, spanning from mitochondrial axonal trafficking to mitochondrial membrane potential and calcium buffering, and including mitochondrial and cytoplasmic viscosity, and cytoplasmic diffusion. By providing a holistic view of the changes occurring in the mouse peripheral nervous system as it ages, we uncover significant differences that display age specificity and are compartment dependent (Fig. 7).

**Figure 7.**
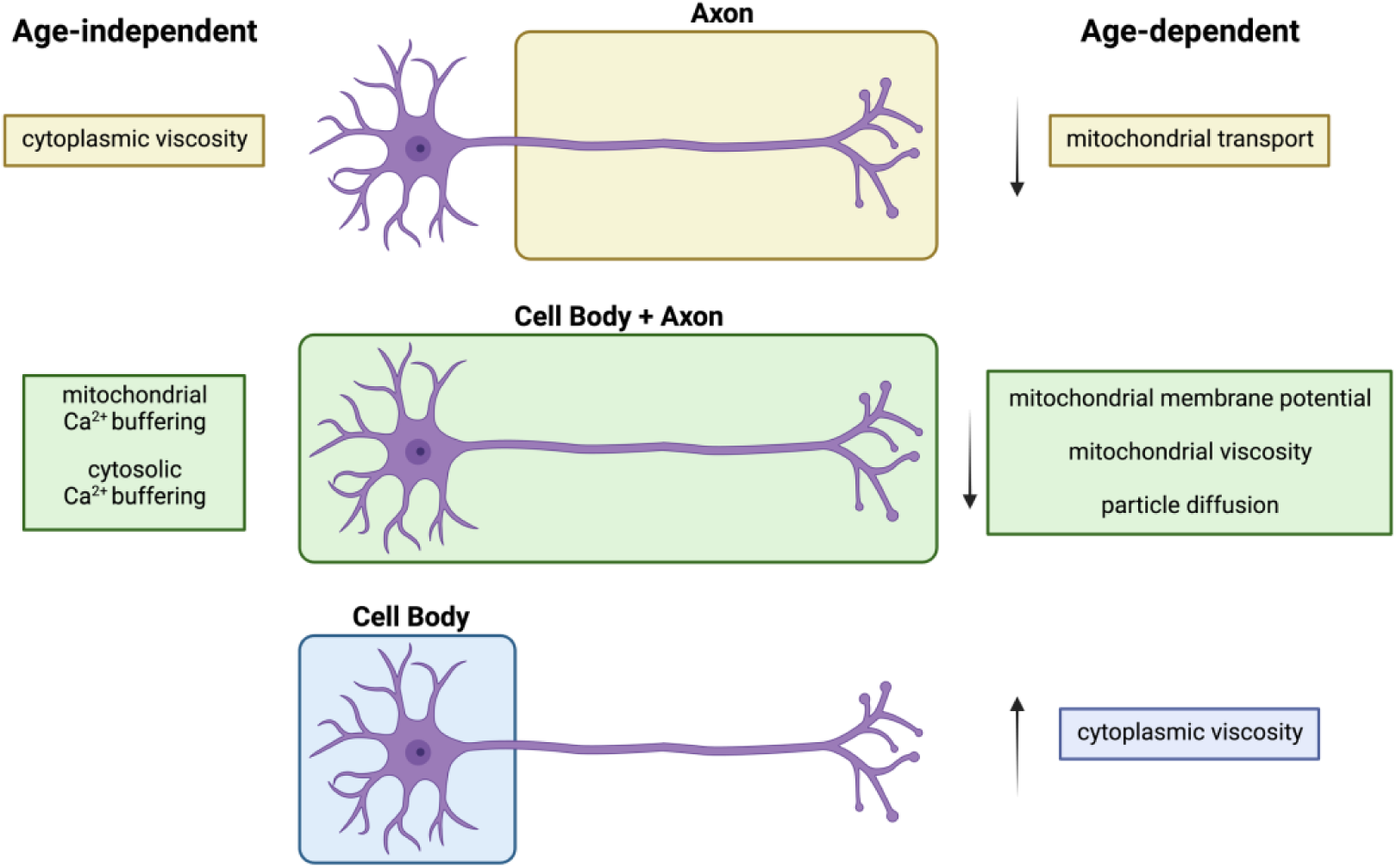
Model of mitochondrial homeostasis and cytoplasmic viscosity during ageing. Decreased axonal mitochondrial transport characterises the ageing process of mouse DRG neurons *in vivo* and *in vitro* (‘Age-dependent’, yellow). The overall decrease in mitochondrial membrane potential observed in aged DRG cultured neurons correlates with reduced intra-mitochondrial viscosity both in the cell bodies and axons (‘Age-dependent’, green) but not with their capacity to buffer calcium, which is unaffected in aged cultures (‘Age-independent’, green). Cytoplasmic viscosity is upregulated in the cell bodies (‘Age-dependent’, blue), but not in axons (‘Age-independent’, yellow). Compartment-dependent and age-specific changes may reflect functional specialisation of the mitochondria and specific regulation of viscosity across sub-cellular compartments and time.

### Mitochondrial axonal trafficking

Altered mitochondrial axonal trafficking is an established feature of ageing peripheral neurons in *Drosophila* and *C. elegans in vivo* (Morsci et al., 2016; Vagnoni et al., 2016) and of mouse sciatic nerve *ex vivo* (Milde et al., 2015). We now show that the number of motile axonal mitochondria declines both in aged DRG neurons *in vitro* and in the sciatic nerve *in situ* (Fig. 7), although the underpinning mechanisms are likely to be different. Firstly, mitochondria travel at significantly different speeds between *in vitro* and *in vivo* systems. Secondly, the total number of axonal mitochondria remains stable *in vitro* showing that decreased transport may not be sufficient to determine the steady-state distribution of mitochondria in axons, as previously suggested (Milde et al., 2015; Vagnoni et al., 2016). On the other hand, decreased mitochondrial motility in the sciatic nerve of older mice may be coupled to impairment in biogenesis or excessive mitochondrial clearance that could lead to an overall reduced number of axonal mitochondria. Our result that mitochondrial area increases in aged axons both *in vitro* and *in vivo* lend support to the notion that altered mitochondrial biogenesis and/or fission and fusion may be a conserved and key feature of neuronal ageing. At the same time, the strength of our comparative study underlies that the DRG *in vitro* system is not a like-for-like replacement of the sciatic nerve *in vivo*: specific mechanisms can differentially affect mitochondrial dynamics *in vitro* and *in vivo* to determine overall mitochondrial distribution.

Decreased mitochondrial motility is accompanied by a surprising increase in the processivity of anterograde mitochondria in aged neurons *in vivo*. It is possible this is a compensatory mechanism triggered by age, similar to upregulation of cargo trafficking observed in response to peripheral nerve injury (Mar et al., 2014; Milde et al., 2015). This result is interesting as it shows that the decline in mitochondrial motility during ageing can be partly reversed in mammalian neurons, as demonstrated previously in ageing *Drosophila* neurons (Vagnoni & Bullock, 2018). Understanding the mechanisms behind the transport upregulation in older neurons might therefore be of therapeutic interest for age-dependent neurodegeneration when decline in cargo trafficking can play an important role in disease progression (Berth & Lloyd, 2023; Sleigh et al., 2019).

### Mitochondrial membrane potential and ion homeostasis

Decline in mitochondrial functionality is an active focus of research in the ageing field (Sharma et al., 2019; Sun et al., 2016) with mitochondrial oxidative stress (Stefanatos & Sanz, 2018) and mitophagy (Caponio et al., 2022; Markaki et al., 2023; Rappe & McWilliams, 2022) thought to play a potentially critical role during ageing. The observation that the ΔΨ_m_ decreases in aged DRG neurons (Fig. 7) implies that the mitochondria become overall less functional and is consistent with reduced mitochondrial bioenergetics observed in iNeurons derived from old human donors (Kim et al., 2018). Upregulation of retrograde transport of dysfunctional mitochondria with reduced ΔΨ_m_ was observed in murine hippocampal (Lin et al., 2017) and chicken DRG neurons (Miller & Sheetz, 2004) after acute mitochondrial stress, suggesting this could be a mechanism for recycling of dysfunctional mitochondria in the cell soma. We now show that this potential safeguarding mechanism may be impaired during ageing due to a strong reduction in mitochondrial motility in both anterograde and retrograde directions. Interestingly, we find that the mitochondria that reside in the cell body are much more polarised than axonal mitochondria and this is observed independently of age. These results suggest that single axonal mitochondria may be metabolically less active than the those found in the interconnected mitochondrial network within the cell body. Alternatively, axonal mitochondria might be more prone to damage, possibly due to a reduced turnover in the axon, where basal mitophagy may occur at different rates than in the cell body (Sung et al., 2016), or to impaired fission and fusion dynamics as a result of decreased trafficking. Whatever the case, our observations support the hypothesis of a mitochondrial functional gradient along the axons whereby mitochondria become progressively less active as the distance from the cell body increases (Baranov et al., 2019; Hagemann et al., 2022). In future studies, it would be interesting to use microfluidics chambers, for example similar to the ones we introduced in this work, to apply stressors to the proximal and distal compartments and observe if mitochondrial transport is differently affected in young and old neurons.

The calcium content of mitochondria has been correlated with their membrane potential (Brini et al., 1999; Duchen, 2000) and motile behaviour (Chang et al., 2011; Niescier et al., 2018), and acute increase in cytoplasmic calcium levels can suppress mitochondrial motility (MacAskill et al., 2009; Wang & Schwarz, 2009). Interestingly, we find that even though mitochondria are overall less polarised in aged DRG cultures when their motility declines, this does not affect their capacity to buffer calcium or the wider neuronal excitability in this context (Fig. 7). Thus, it is conceivable that the population of healthy mitochondria in the older DRG neurons is sufficient to compensate for the dysfunctional organelles, although we cannot exclude that the calcium uptake capability is retained in depolarised organelles.

### Mitochondrial viscosity

Reduced ΔΨ_m_ is accompanied by an overall reduction in the intramitochondrial viscosity in aged DRG (Fig. 7), which could conceivably reflect an adaptation to metabolic activity (Persson et al., 2020; Puchkov, 2013) that becomes altered with age. Alternatively, changes in mitochondrial membrane fluidity as a result of oxidative damage (Eckmann et al., 2014; Mecocci et al., 1997) or intra-mitochondria molecular dilution due to enlarged area (Fig. 1-2) could explain reduced viscosity. Interestingly, mitochondria of human fibroblasts that are deficient for complex I activity display enhanced protein diffusion (Koopman et al., 2008), likely a sign of reduced viscosity, while mitochondria hyperpolarisation correlates with increased viscosity in HepG2 cells (Bulthuis et al., 2023). Thus, our data support the idea that the reduced ΔΨ_m_ and mitochondrial viscosity in aged DRGs are mechanistically linked.

Contrarily to ΔΨ_m_, however, mitochondrial viscosity across compartments remained uniform, which suggests independent mechanisms might regulate ΔΨ_m_ and viscosity in different compartments. Mitochondrial viscosity was shown to be selectively unaffected by acute osmotic stress compared to other organelles (Chambers et al., 2018). Therefore, it is possible that mitochondrial viscosity is less sensitive to stressors that could determine a difference in ΔΨ_m_ across compartments. Overall, our study adds significantly to the notion of mitochondrial functional specialisation that has been observed across different neuronal populations (Fecher et al., 2019) and within subcellular compartments (Faitg et al., 2021). Future studies should be aimed at further characterising the mitochondrial microviscosity environment to assess whether sub-mitochondrial domains are differently affected by age in different cellular compartments.

### Cytoplasmic viscosity

Whether cytoplasmic viscosity and diffusion undergo significant changes during ageing that can affect intracellular trafficking is not known. Previous studies showed that increased viscosity reduces both endosomal trafficking and microtubule dynamics (Molines et al., 2022; Nunes et al., 2015). Changes in cytoplasmic viscosity are thought to depend on alterations of molecular crowding, with likely repercussions on wider cellular functionality (Leduc et al., 2012; Metzner et al., 2022; Sabharwal & Koushika, 2019; Sood et al., 2018). Using TR-FAIM, we find that axonal cytoplasmic viscosity does not change with age and so it is unlikely to have a major impact on the biology of the axon, including the decrease in mitochondrial motility that we observe in older DRG neurons (Fig. 7). However, we find that, contrarily to axons, the viscosity of the cell bodies increases in aged cultures. Why cell bodies selectively display increased viscosity is not clear. This could result from an imbalance in protein translation (Delarue et al., 2018), a potential temperature difference across compartments as the cell ages (Persson et al., 2020), for example as a consequence of reduced metabolic activity (Parry et al., 2014), or be the product of compartment-dependent difference in plasma membrane stiffness (Kubánková et al., 2019; Lamoureux et al., 2010; Sugawa et al., 1996). Whatever the case, we demonstrated that acutely increasing cell body viscosity using molecular crowders is sufficient to strongly reduce mitochondrial export into the axons and overall transport rates. We are aware of the challenge posed by recapitulating age-dependent processes with acute, albeit well controlled, experimental treatments. Nevertheless, our results are strongly suggestive of a scenario in which the age-dependent increase in cell body cytoplasmic viscosity contributes to reducing axonal transport rates, with a likely wider impact on cellular functionality in aged cells. The mechanism behind this process is not known. Our observation that mitochondria appeared more clustered and less dynamics in the DRG cell bodies after PEG and sorbitol treatments (Movie 6), suggests this might be a likely cause for the decreased mitochondrial export into the axons. However, it would be valuable to establish in future studies if viscosity alters the molecular composition of the AIS-like filter zone in cultured DRGs (Farías et al., 2015; Gumy et al., 2017) to determine the abundance of selected cargoes entering the axons in ageing cells.

The apparent reduction of mitochondrial axonal transport speed after sorbitol treatment of the cell bodies is interesting: it suggests that, in the highly crowded environment of the cell body, the mitochondria and/or the mitochondrial transport machinery may undergo molecular changes that affect their long-term trafficking. At first, this result would not seem consistent with the unchanged mitochondrial velocity measured in the axons of aged, cultured neurons (Fig. 1H-I). However, we think that the acute increase in viscosity reveals that reduced speeds may be the precursor to overall reduced mitochondrial motility observed during ageing (Fig. 1C), when exposure to increased viscosity is likely to occur more gradually.

These data further support the hypothesis that failed mitochondrial export from the soma is a primary contributor of reduced axonal trafficking during ageing. In turn, reduced mitochondrial supply may trigger an imbalance in mitochondrial fission and fusion in axons leading to enlarged organelles. It will be intriguing to test the validity of our model in future work by studying if manipulating cell body viscosity directly affects a pool of predominantly axonal mitochondria in aging neurons.

### Molecular diffusion

We find that the molecular diffusion in the neuronal cytoplasm, ascertained using the GEM probe, decreases with age in both soma and axons (Fig. 7). While this is consistent with the increased viscosity of the cell soma, it seems apparently at odds with the constant axonal viscosity measured by TR-FAIM. However, our data are in accordance with the notion that macromolecular crowding in complex cellular environments can cause deviation from the Stokes-Einstein law that relates diffusion and viscosity (Zustiak et al., 2011). Moreover, TR-FAIM measures rotational diffusion of a dye whereas GEMs explore lateral diffusion in the mesoscale. Therefore, we speculate that older axons may develop proteinaceous aggregates and/or molecular condensates that, while not sufficient to alter overall viscosity, can provide significant spatial hindrance by increasing local crowding in small calibre DRG axons. This could explain both the slower diffusion of the 40 nm GEM particles and the well documented decrease in organelle motility observed during ageing and neurodegeneration. In future studies, it would be valuable to determine whether local viscosity and diffusion in proximal and distal axonal regions display different rates, and to establish if this is a general feature of the nervous system or confined to a specific subset of neurons. Ultimately, it would be relevant to understand whether age-dependent neurodegenerative diseases are characterised by specific viscosity and diffusion patterns and whether this could be exploited to model disease progression.

Overall, this work will provide an important reference for future studies aimed at better understanding mitochondrial functionality and the role of the cytoplasmic milieu in cellular homeostasis.

## ACKNOWLEDGEMENTS

We thank personnel of the Denny Brown Laboratory for assistance with mouse colonies, Gábor Mórotz for suggestions regarding calcium imaging experimental setup, BFK Lab for assistance with MSD analyses, the Wohl Cellular Imaging Centre at King’s College London for help with light microscopy, Frank Hirth and members of the Vagnoni lab for comments on the draft manuscript. Fig. 1A, 2A, 6A and 7 were created using https://www.biorender.com. This project was funded by a NC3Rs David Sainsbury Fellowship NC/N001753/2 and SKT grant NC/T001224/1 (to A.V.), a BBSRC CTP award BB/T508597/1 (to A.V.), Royal Society Research Grant RGS\R2\180450 (to A.V.), an Academy of Medical Sciences Springboard Award SBF004/1088 (to A.V.), a van Geest Fellowship in Dementia and Neurodegeneration and van Geest PhD studentship awards (to A.V.), a King’s Together Multi and Interdisciplinary Research Scheme Wellcome Trust Institutional Strategic Support Fund 204823/Z/16/Z (to A.V.), an European Union’s Horizon 2020 research and innovation programme, grant agreement No. 857524 (to A.V.), and by the Alzheimer’s Research UK London Network Centre grant ARUK-NC2020-KCL, a BBSRC award BB/R004803/1 (to K.S.), Medical Research Council awards MR/S006990/1 and MR/Y010949/1 (to J.N.S.), the UCL Neurogenetic Therapies Programme funded by The Sigrid Rausing Trust (to J.N.S., G.S.), a Wellcome Trust Sir Henry Wellcome Postdoctoral Fellowship 103191/A/13/Z (to J.N.S.), Wellcome Trust awards 107116/Z/15/Z and 223022/Z/21/Z (to G.S.), and a UK Dementia Research Institute Foundation award UKDRI-1005 (to G.S.). D.S. and I.E.S. were supported by a BSCB Studentship and a LIDo iCASE Studentship, respectively.

For the purposes of open access, the author has applied a Creative Commons Attribution (CC BY) licence to any Accepted Author Manuscript version arising from this submission.

The authors declare no conflict of interest.

## MATERIALS AND METHODS

### Study approval

Experimentation involving mice was performed under license from the UK Home Office in accordance with the Animals (Scientific Procedures) Act (1986). Work conducted at University College London was approved by the Queen Square Institute of Neurology Ethical Review Committee and experiments performed at King’s College London were approved by King’s College Denmark Hill Ethical Review Body.

### Animals

Mice were maintained under a 12 h light/dark cycle at constant room temperature (≈21°C) with free access to water and food. Cages were enriched with nesting material, plastic/cardboard tubes and wooden chew sticks. MitoMice (B6.Cg-Tg(Thy1-CFP/COX8A)S2Lich/J – RRID:IMSR_JAX:007967) were obtained from The Jackson Laboratory and maintained as hemizygous × wild-type breeding pairs on a C57BL/6J background. Presence of the CFP transgene was confirmed by a standard condition polymerase chain reaction using forward primer 5’-GAC GTA AAC GGC CAC AAG TT-3’ and reverse primer 5’-GTC CTC CTT GAA GTC GAT GC-3’. C57BL/6J wild-type mice were obtained from Charles River Laboratories UK Ltd. Mice of both sexes were used in all experiments. DRGs obtained from MitoMice were used for live imaging of mitochondrial transport in mass cultures and Ca^2+^ imaging experiments, while DRGs from C57BL/6J mice were used for live imaging of mitochondrial membrane potential and mitochondrial viscosity, TR-FAIM and microrheology experiments, and mitochondrial transport in microfluidic chambers. Young and old cells were imaged in the same or in contemporary experimental sessions, whenever possible, rather than 20 months apart.

### DRG dissection and mass culture

DRGs were dissected from the spinal cord of euthanised adult mice (Sleigh et al., 2016). Approximately 10 DRG pairs (*i.e.*, 20 DRG) were collected for each mouse, and kept on ice in Ca^2+^/Mg^2+^-free HBSS with 1% Pen/Strep until dissociation. DRGs were cultured as previously described (Sleigh et al., 2017b); briefly, after removing the HBSS, the DRGs were incubated in 4 mg/ml collagenase type II (Thermo Fischer Scientific #17101015) and 4.7 mg/ml dispase type II (Sigma-Aldrich #D4693) in Ca^2+^/Mg^2+^-free HBSS at 37°C for 10-15 min with periodic gentle agitation. The collagenase/dispase solution was replaced with pre-warmed F12 medium (F12 (Gibco) containing 10% heat-inactivated-FBS (hi-FBS) and 1% Pen/Strep (both from Thermo Fischer Scientific)). Upon centrifugation, cells were resuspended in F12 medium and triturated using fire-polished glass Pasteur pipettes. After centrifugation at 200 *g* for 5 min, cells were resuspended in 320 μl of fresh DRG medium (F12, 10% hi-FBS, 1% Pen/Strep, 20 ng/ml mouse GDNF (Peprotech #450-44), 50 ng/ml mouse NGF (Peprotech #450-34), 20 ng/ml human NT3 (Peprotech #450-03)) and plated as single 20 μl drops (1-2 drops/well or coverslip) at 37°C, 5% CO_2_ for 1-2 h. 0.3-1 ml of DRG medium was then gently added to each well and cells were returned to the incubator. Cells were plated in μ-Slide 8-well glass bottom imaging chambers (Ibidi #80827), in 8- and 4-well Nunc Lab-Tek chambered coverglass (Thermo Fischer Scientific) or 18-mm round glass coverslips (Marienfeld), depending on the experiments. 20 μg/ml laminin (Corning #354232) in Ca^2+^/Mg^2+^-free HBSS (Gibco) was used to coat the culture surface for at least 2 h before plating.

### Microfluidic chambers (MFCs) preparation and DRG cultures

A preparation of 10:1 polydimethylsiloxane (PDMS) and curing agent (Sylgard 184 Silicone Elastomer Kit) was vigorously mixed (approximately 3.5 ml per MFC mould), then bubbles removed in a desiccator vacuum pump for 30 min. PDMS mix was carefully poured into 500 μm MFCs moulds, then baked at 65°C for 60-75 min. Baked PDMS was removed from moulds and punched to create chips. Glass-bottomed dishes were coated with poly-D-lysine for 3-16 h to create static attraction to the PDMS chips, washed 5 times with sterile H_2_O, then PDMS chips were stuck down. The channels were treated with 0.8% BSA overnight, then coated with laminin as described previously (Rhymes et al., 2022).

DRGs were isolated as described above and resuspended in 40 µl of DRG medium. 10 µl of cell suspension were added to one well of the MFC cell body compartment (cell body well 1, Fig. S7A) held at a slightly tilted angle to aid distribution throughout the cell body channel (Fig. S7A), while 20 µl of DRG medium were added to the corresponding axonal well (axon well 1, Fig. S7A). Cell suspension was incubated for approximately 1 h at 37°C before adding 140 µl of DRG medium dropwise across both wells of the cell body compartment (cell body well 1 and 2, Fig. S7A), and 80 µl across the wells of the axonal compartment (axon well 1 and 2, Fig. S7A).

### Live imaging and analysis of mitochondrial transport and morphology in mass cultures of DRG neurons

DRGs were imaged at 3-4 days *in vitro* (DIV). Before imaging, cells were incubated with 200 nM of MitoTracker Deep Red for 30 min and the dye washed off prior to imaging. Mitochondrial axonal transport was recorded with a Nikon spinning disk system comprising of a CSU-X1 scanning head (Yokogawa) and a Nikon Eclipse Ti-E inverted microscope equipped with a Du 897 iXon Ultra electron-multiplying charge coupled device camera (Andor) and a 60× CFI Apo/1.4 NA objective. Mitochondrial movements were typically recorded for 2 min with an acquisition rate of 0.5 frames/s, 640 nm excitation laser (5-10% power), 100-150 ms exposure, gain 1, EM gain 100. Temperature and CO_2_ were maintained at 37°C and 5%, respectively, via a temperature unit with CO_2_ control and a microscope enclosure (Okolab). Axonal transport was recorded approximately 100 μm away from the cell body and kymographs were produced by first straightening 100 μm of axonal tract (approximately in mid axon) using the Straighten plugin and followed by the Velocity Measurement Tool in ImageJ, as described previously (Annuario et al., 2022; Mattedi et al., 2023). Mitochondrial speed, run length and number of total mitochondria were analysed directly from the kymographs. Time-lapse movies were used to unambiguously confirm the mitochondrial tracks, when necessary. A mitochondrion was defined as motile if containing at least one continuous bout of net motion of at least 2 μm (a ‘run’) in the anterograde or retrograde direction. Because mitochondrial traces can be composed of multiple runs of variable speeds, velocity is defined as the average run velocities per single mitochondrion. Mitochondrial displacement is defined as the sum of the mitochondrial run lengths per single mitochondrion. Bidirectional mitochondria display at least one run in both the anterograde and retrograde directions.

Mitochondrial morphology was quantified by calculating single mitochondrial surface area, length, and elongation, using the NIS Elements Analysis software. Mitochondrial length was defined as the length of the medial axis of the mitochondria while elongation was defined as ratio between maximal and minimal Feret’s diameter. The outline of each mitochondrion was obtained by manually adjusting the intensity threshold for each movie and quantification of morphological parameters was performed on individual representative timeframes at which mitochondria were best visible. The same movies were utilised to quantify mitochondrial transport and morphology.

### DRG treatments and live imaging in MFCs

DRGs were imaged at 3-4 DIV. Prior to imaging, the cell body compartment was treated with either PEG (polyethylene glycol, MW 8,000, Sigma) for 1 h or with sorbitol (Sigma) for 30 min. PEG and sorbitol were separately diluted in DRG medium to 20% and 300 mM, respectively, and 100 µl of each treatment were added dropwise across the cell body wells. 150 µl of DRG medium without treatments were added across the axonal wells. Following the initial incubation with the crowding agents, the medium in the cell body wells was replaced with 100 µl DRG medium with treatments supplemented with 200 nM MitoTracker Green and 1X CellMask Orange (CMO, ThermoFisher). The medium in the axon wells was replaced with 150 µl medium containing 1X CMO only. After 30 min, the dyes were washed off and the neurons imaged immediately, with the treatments kept on in the cell body compartments throughout imaging.

Mitochondrial axonal transport was recorded in the initial portion of the microgrooves containing axonal projections using the same Nikon spinning disk system used for the mass cultures and a 60× CFI Apo/1.4 NA objective. Mitochondrial movements were recorded for 1 min with an acquisition rate of 0.5 frames/s, 488 nm excitation laser (10% power), 200 ms exposure, gain 1, EM gain 100. Mitochondrial flux analysis was performed by counting how many times in 1 min the mitochondria crossed a line drawn in the initial segment of the microgrooves, close to the somal compartment.

Mitochondrial mass was defined as the total mitochondrial surface area in a region of interest of 54 µm adjacent to the somal compartment and quantified as described above for DRG cultures.

Cell body area was calculated by drawing a line around the border of the cell body, as defined by CMO staining, using the Fiji Polygon selection tool.

Low magnification images to capture the length of the MCF microgrooves were acquired using a 20×/0.75 NA CFI Plan Apochromat VC air objective, and the ‘Large Image’ function on the NIS Acquisition software.

### Live imaging and analysis of mitochondrial transport and morphology in the mouse sciatic nerve

Exposure and *in vivo* imaging of intact sciatic nerve axons in anaesthetised mice were performed under terminal anaesthesia as previously described (Sleigh et al., 2020; Tosolini et al., 2021). Time-lapse imaging of mitochondrial transport was executed using an inverted LSM780 laser-scanning microscope (ZEISS) and 63× Plan Apochromat oil immersion objective lens (ZEISS) at 100× digital zoom. Images were acquired at a rate of 0.5 frames/s for 8 min, with laser power 2%, resolution 1024×1024 and line averaging of 4. Analysis of mitochondrial motility was carried out essentially as previously described (Vagnoni & Bullock, 2018). Tracking of mitochondria was performed manually in Fiji/ImageJ on the raw movies using MtrackJ by marking the start and end of each run and tracking was performed blind to the age and sex of the animals. Speed and displacement were calculated from MtrackJ-generated tracks of runs ≥ 2 μm, as defined above. Total mitochondrial number was averaged from two frames at the beginning and at the end of the time series. The StackReg plugin of Fiji/ImageJ was used to correct for sample drifting, if necessary.

Mitochondrial morphology was quantified as described above for DRG cultures.

### Live imaging of mitochondrial membrane potential and siRNA treatment

DRGs were cultured to DIV3-4 and incubated with 100 nM MitoTracker Green FM (Thermo Fischer Scientific) in DRG medium for 30 min at 37°C. After washing off the dye, 50 nM TMRM (Thermo Fischer Scientific) was added to fresh medium and incubated for 15 min. After washing off the TMRM dye with fresh medium, imaging was performed with a Nikon A1RHD inverted confocal microscope, equipped with GaAsP and multi-alkali PMT detectors and a 60×/1.4NA Plan Apochromat oil immersion objective. Cells were kept at a constant temperature of 37°C, 5% CO_2_ throughout imaging. For both 488 nm (MitoTracker Green) and 561 nm (TMRM) excitation (laser power of 1% and 0.3%, respectively), we set the pixel dwell time at 2.2 μs, image size at 512×512, averaging at 2× and pinhole size at 39.6 μm. Low dye concentration and laser power were used to prevent TMRM quenching and FRET between dyes (Mitra & Lippincott-Schwartz, 2010). Quantification of fluorescent signal was performed using the raw intensity values in Fiji/ImageJ. TMRM fluorescence intensity was measured and divided by the fluorescence signal of MitoTracker Green to obtain the normalised value of the mitochondrial membrane potential. For presentation purposes, the ‘Smooth’ filter in Fiji/ImageJ was used to reduce salt and pepper noise.

Dharmacon Accell SMARTpool Trak1 siRNA (Cat#E-056081-00-0010) and non-targeting control siRNA (Cat#D-001910-10-05) from Horizon Discovery were reconstituted to 100 μM following the manufacturer’s instruction. When cultured DRGs reached DIV1, siRNAs were mixed with the Accell Delivery Media (ADM, Cat #B-005000-100) and, after removing 200 μl of conditioned medium from each well of an 8-well imaging chamber, 200 μl of siRNA-ADM complexes were delivered dropwise on the cells at 1 μM final concentration (300 μl total medium volume/well). After 72 h (DIV4), neurons were incubated with 4 μM of the JC-1 dye (Thermo Fisher Scientific) for 30 min and the medium was replaced before imaging. The aggregation state of the dye is affected by ΔΨ_m_ with the fluorescence emission shifting from green to red in more polarised mitochondria. The fluorescence emission at ∼ 590 nm (red) and ∼ 530 nm (green) was captured with emission filters ET595/50m and ET525/50m (both from Chroma Technology Corporation), respectively. The ∼ 590/530 fluorescence ratio gives a measurement of ΔΨ_m_. For trafficking studies using MitoTracker DeepRed, 0.5 μl non-targeting Green/6-FAM siRNA (Cat#D-001950-01-05) was added to the siRNA-ADM complexes to assess the efficiency of siRNA delivery.

### Live cell calcium imaging

DRGs were imaged with a Nikon Ti-E inverted epifluorescence microscope equipped with a Nikon 20×/0.8 NA Plan Apochromat air objective, a mercury lamp (Nikon Intensilight C-HGFI) for illumination and a Dual Andor Neo sCMOS camera for detection. Cells were plated on laminin-coated 18 mm coverslips and treated with 2 μm Rhod2-AM and 4 μm Fluo4-AM (both from Thermo Fisher Scientific) for 15 min at 37°C in external solution (145 mM NaCl, 2 mM KCl, 5 mM NaHCO_3_, 1 mM MgCl_2_, 2.5 mM CaCl_2_, 10 mM glucose, 10 mM HEPES pH 7.4) (De vos et al., 2012) prior to imaging. Cells were imaged at DIV4 in a Ludin imaging chamber (Type 1, Life Imaging Services) in external solution at 1 frames/s for 3 min using GFP/mCherry filter set and exposure time of 30 ms. Cells were imaged for 30 s before transiently perfusing the imaging chamber with 50 mM KCl with a peristaltic pump (Ismatec). KCl was washed off after 1 min and each coverslip was subjected to only one round of perfusion and imaging.

To quantify KCl-induced Fluo4 and Rhod2 response, a region of interest (ROI) was drawn to outline single cell bodies and axonal tracts from each field of view and the fluorescence intensity of the dyes measured in Fiji/ImageJ from raw images displaying non-saturated fluorescent intensity. Response curves were aligned at the base of the response peak and the fluorescence intensity at every time point was normalised to the value of the first frame. Fluorescence ‘peak’ was defined as the maximum fold-change value reached during imaging and the time to reach the peak was calculated as the Δt between the ‘peak’ timepoint and the time of first response to stimulation, as reported previously (Mattedi et al., 2023). Time to decay was defined as the time necessary for the fluorescence signal to decline by 80% of the peak value.

### GEMs viral preparation and transduction

To obtain GEMs-encoding lentiviral particles, the pCMV-PfV-Sapphire-IRES-tdTomato plasmid (Addgene, #116934) (Delarue et al., 2018) was transfected into Lenti-X 293T cells (TaKaRa, #632180) using the polyethylenimine(PEI)-based Transporter 5 transfection reagent (Polysciences, #26008) and a 2^nd^ generation viral packaging system comprising gag/pol/tat/rev (psPAX2, Addgene, #12260) and VSV-G (pMD2.G, Addgene, #12259). Cells were seeded in a 15 cm dish in 22.5 ml medium (DMEM with 4.5g/l glucose and pyruvate, 10% hi-FBS, 1x GlutaMAX) at 37°C with 5% CO_2_ until they reached 60-80% confluency. The medium was then replaced and cells incubated for 2 h with fresh medium containing 2% FBS. DNA-PEI complexes (14.6 μg psPAX2, 7.9 μg pMD2.G, 22.5 μg GEM-transfer vector and 135 μl Transporter 5) were prepared according to the manufacturer’s instructions and added dropwise to the cells. After overnight incubation, the medium was replaced with 17 ml fresh medium with 2% FBS, the supernatant harvested 3 times every 8-12 h and kept at 4°C. Typically, the three harvests were centrifuged at 500 *g* for 5 min at 4°C before filtering through 0.22 μm filter units and further ultracentrifugation at 50,000 x *g* for 2 h at 4°C. After gently discarding the supernatant, the viral pellet was let dry for 2-3 min before resuspension in DMEM without FBS or Opti-MEM to obtain a 1000-fold concentration and stored at -80°C in 10-20 μl aliquots. Between 1-10 μl of concentrated virus was added to each well of a μ-Slide 8-well glass bottom imaging chambers.

### GEMs imaging and single particle analysis

DRGs were plated in μ-Slide 8-well glass bottom imaging chambers and the concentrated lentiviral particles added at DIV4. The medium was replaced after 24 h with fresh medium without growth factors and the cells imaged after an additional 36 h (DIV7) by spinning disk confocal microscopy with a Nikon 100×/NA 1.40 CFI Plan Apochromat VC oil immersion objective and 100 ms continuous acquisition for 30 s, 488 nm laser at 40-50% intensity, gain 1, EM gain 250. Cells expressing similar levels of transduced construct, as assessed by the tdTomato signal, were selected for imaging.

Single particle tracking (SPT) was performed on raw time series using TrackMate on selected ROIs defining the cell body and axonal compartments using the DoG detector for small spot detection (<5 pixels), with estimated particle diameter set at 0.4-0.5, thresholds between 0.4-0.6. with median filter and sub-pixel localization. The Simple LAP tracker was used to link the particles throughout the movie (linking max distance = 1, gap-closing max distance = 1.6, gap-closing max frame gap = 2). Tracks were then exported and analysed in MATLAB. Time-averaged mean square displacement (MSD) was calculated via creation of a SPT Analyser using a custom script (available on request). The mean diffusion coefficient (*D_eff_*) was calculated by fitting the first 10 s of the time-averaged MSD with the equation *MSD (t) = 4Dt^α^*, where α is the exponent describing the anomalous diffusion and *D* is the diffusion coefficient.

### FLIM

Calibration and FLIM measurements were carried out essentially as in Steinmark et al. (Steinmark et al., 2019). DRGs were plated in μ-Slide 8-well glass bottom imaging chambers. On the day of imaging (DIV3-4), cells were washed in external solution and incubated with 1 μM FMR-1 for 30 min at 37°C and the dye was washed off prior to imaging. Cells were imaged on an inverted TCS-SP2 microscope (Leica Microsystems) using two SPC-150 TCSPC boards (Becker & Hickl). FMR-1 was excited using a picosecond-pulsed (90 ps optical pulse width, 20 MHz repetition rate) 467 nm diode laser (Hamamatsu, Japan) using a HCX PL APO CS 63X 1.2 NA water objective, with a 485 nm dichroic mirror and a 514/30 nm band pass filter. All measurements were carried out at 37°C and 5% CO_2._ Analyses were performed at single cell level by measuring the fluorescence lifetime separately in cell bodies and proximal-mid axons regions.

### TR-FAIM

DRG neurons were grown to DIV3-4, as for the FLIM experiments, and labelled with 1 µM BCECF (Themo Fisher Scientific #B1150) for 30 min at 37°C. The dye was washed off prior to imaging in FluoroBrite DMEM. Time-resolved anisotropy measurements were conducted on the same TCS-SP2 microscope system used for FLIM, with the BCECF dye excited using a picosecond-pulsed laser. In this case, the fluorescence emission was split into its parallel and perpendicular components using a polarizing beam splitter (Edmund Optics).

#### Calibration curve

Rotational correlation times of BCECF in solutions of increasing viscosity (through addition of glycerol) were measured and plotted. To avoid depolarisation effects from the objective, a 5x NA 0.15 objective was used and measurements were conducted at room temperature, but otherwise the setup was identical to that described above. The anisotropy was calculated using the equation:

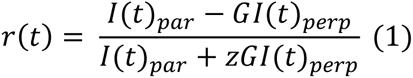

Where z = 2 and G is a correction factor used to correct for the different sensitivity of the two detectors. It is calculated by taking the tail value of:

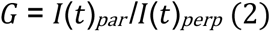

G-values used were around 1 +/-0.1. Here the G-factor was measured using a solution of fluorescein in water and 10% glycerol. The calibration curve was calculated to avoid assumptions about molecular shape and diffusion boundary conditions in a classic Stokes-Einstein-Debye calculation.

#### Fluorescence Anisotropy decay

For each raw neuron image, two regions of interests (ROIs) were extracted: from the axon and the soma. For each ROI, a single set of parallel and perpendicular decays were generated by summing the photon counts in each time bin over all pixels within the given ROI. Anisotropy pre-processing, curve fitting and graphing were performed in OriginLab. The perpendicular (*I_perp_*) and parallel (*I_par_*) decays for each ROI of each image were aligned and the anisotropy decay was calculated using equation (1) where z, due to the high NA of the objective, was determined to be 1, and G was determined as described above. Analyses were performed at single cell level by calculating anisotropy separately on cell bodies and proximal-mid axon region of the same cell. For calculating the anisotropy of an entire cell, the soma and axon photon counts were summed up and analysis was performed as described. It was established that the hindered rotation model (below) provides an optimum fit for most anisotropy decays, accounting for a restricted model of rotation in all environments.

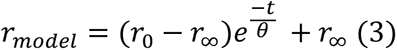

Where r_0_ is the initial anisotropy, r_∞_ is the limiting anisotropy and θ is the rotational correlation time. The weighting factor v(r) for the anisotropy fits was the variance of r (Lidke et al., 2005):

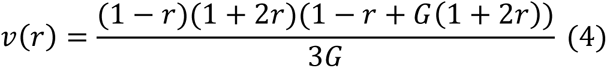

Only the first 10 ns of the decay were used for the fit and only meaningful fits with an acceptable adjusted R-squared value and standard errors of the parameter fits were accepted for further analysis.

### Statistical analyses

Details of the statistical analyses are reported in the figure legends. Shapiro-Wilk test was performed to test data for normal distribution before determining the appropriate statistical test. Sample sizes, which were predetermined using power calculations and previous experience, are also reported in figure legends.

---

**Figure S1.**
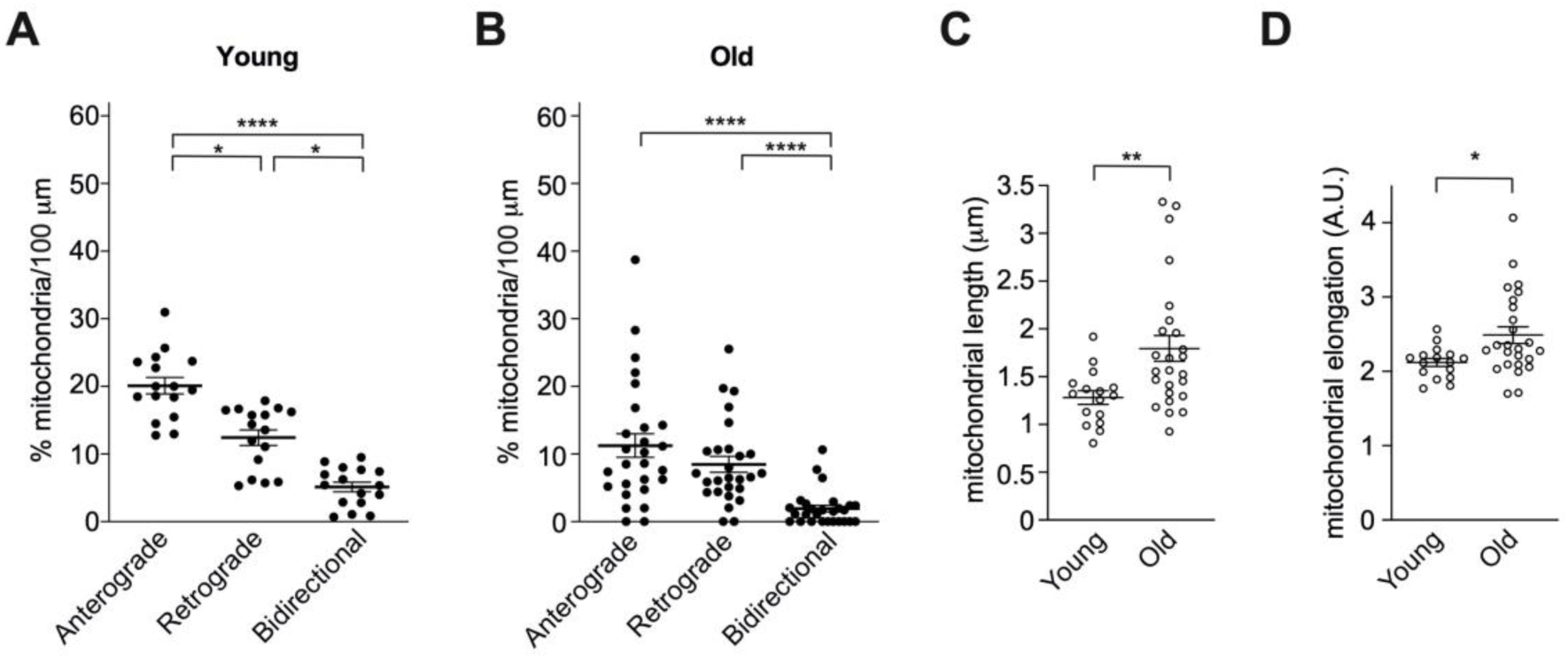
Proportion of motile mitochondria displaying anterograde, retrograde and bidirectional motility in young **(A)** and old **(B)** DRG neurons *in vitro* (relative to Figure 1C). A general decrease of motility is observed during age. **(C-D)** Quantification of mitochondrial length and elongation (see Methods) in young and old cells. Individual data points represent fields of view (FOV) imaged (1-3 axons/FOV). Data are shown as mean ± SEM. Kruskal-Wallis with Dunn’s multiple comparisons test (A-B) and Mann-Whitney test (C-D). * p < 0.05; ** p < 0.01; **** p < 0.0001.

**Figure S2.**
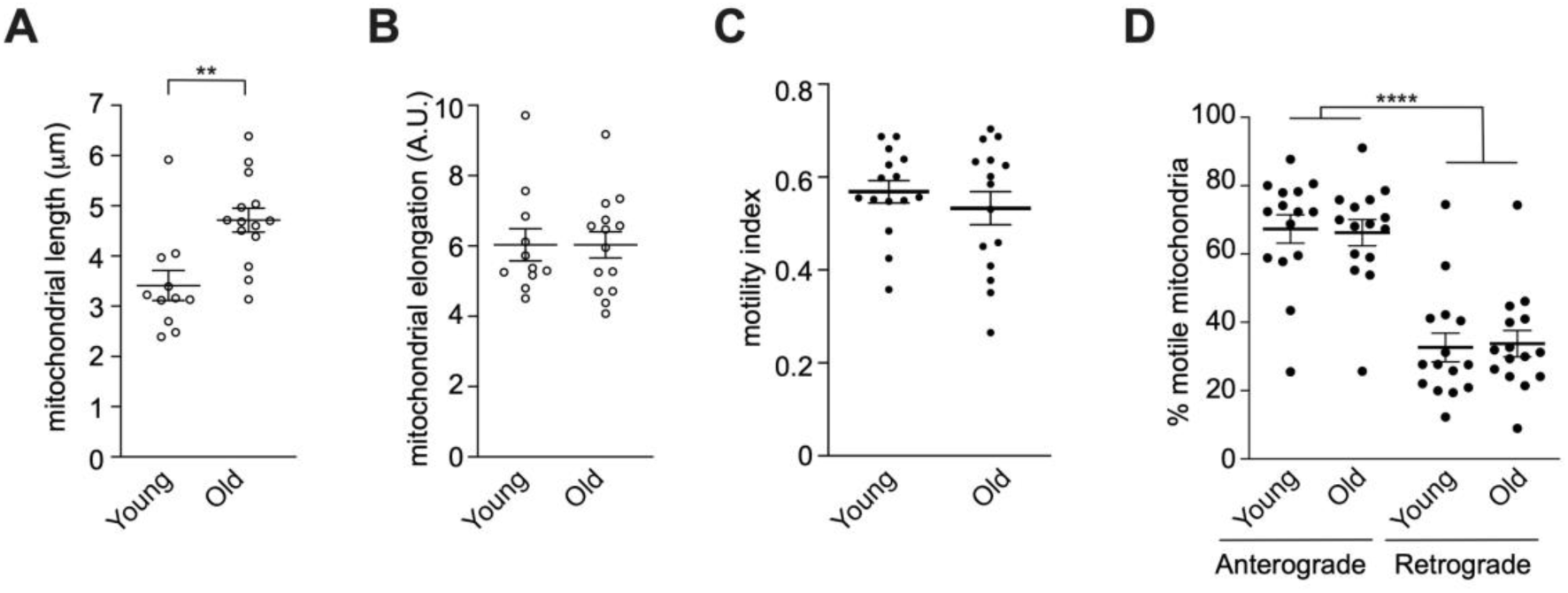
**(A-B)** Quantification of mitochondrial length and elongation (see Methods) in young and old sciatic nerve axons *in vivo*. Individual data points represent sciatic nerve axons. **(C)** There is no difference in the mitochondrial motility index between young and old neurons (*i.e.*, total moving mitochondria/(total moving + stationary mitochondria)). **(D)** The relative proportions of anterograde and retrograde mitochondrial transport are not changed by age in the sciatic nerve, indicating that the reduction of both types of transport contributes to the overall transport decline (Figure 2D). Individual data points represent fields of view (FOV) imaged (1-3 axons/FOV). Data are shown as mean ± SEM. Mann-Whitney test (A-B), unpaired student’s t-test (C), two-way ANOVA with Tukey’s multiple comparisons test (D). ** p < 0.01; **** p < 0.0001.

**Figure S3.**
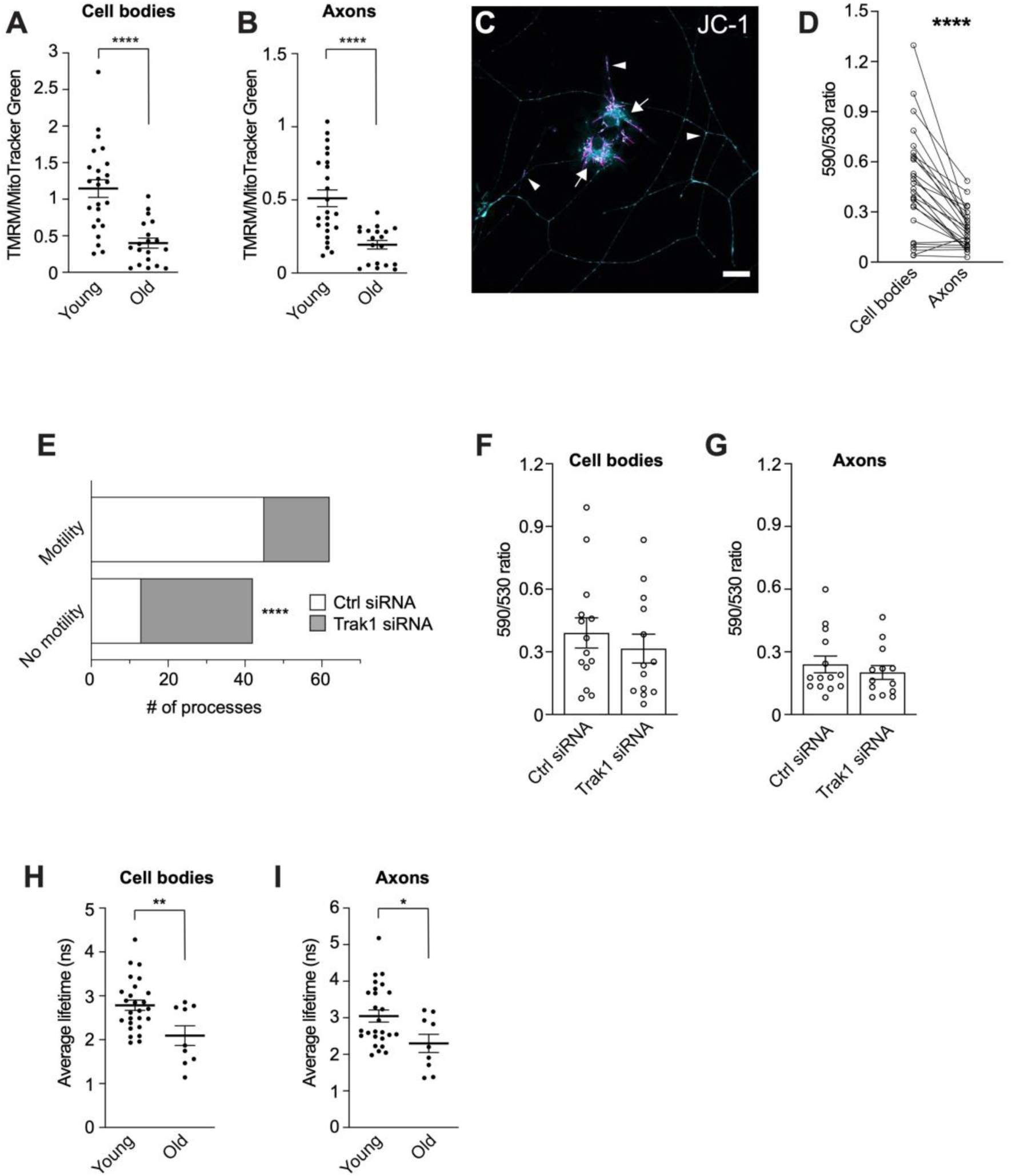
**(A-B)** Quantification of the ratio between TMRM and MTG in neuronal cell bodies (A) and axons (B) indicates that the ΔΨ_m_ declines during ageing in both compartments. **(C)** DRG neurons stained with the JC-1 vital dye and pseudocoloured in cyan and magenta (∼ 530 and ∼ 590 nm emission, respectively). Arrows, somal compartments; arrowheads, axons. Note the higher polarisation of somal mitochondria compared to axonal mitochondria, indicated by widespread magenta, corroborating the data in Fig. 3A-G. Scale bar: 20 μm. **(D)** Quantification of the 590/530 JC-1 ratio from young DRG neurons indicates that the axonal mitochondria display reduced ΔΨ_m_ compared with cell body mitochondria. Data points represent the cell bodies and axons imaged, with each neuron providing both cell body and axons for paired analysis. **(E)** Increased number of DRG neuron axons displaying no motile mitochondria after Trak1 RNAi. A process is scored as displaying motility if at least two mitochondria are motile within 1 min of imaging. Number of axons analysed are 58 (Ctrl siRNA) and 46 (Trak1 siRNA). **(F-G)** Trak1 RNAi does not significantly change the 590/530 JC-1 ratio in the cell bodies (F) or axons (G) of DRG neurons, suggesting the ΔΨ_m_ is not affected by this manipulation. Data points represent individual cell bodies and axons, respectively. Each neuron analysed provides both cell body and axons to the analysis. DRGs for JC-1 quantifications were obtained from 3 young (P102-105) male mice. **(H-I)** Mitochondrial viscosity decreases during age both in the cell bodies (H) and axons (I) of DRG neurons. Data points represent cell bodies and axons imaged. Data are shown as mean ± SEM. Unpaired student’s t-test (A, F, H), Mann-Whitney test (B, G, I), paired student’s t-test (D), Fisher’s exact test (E). * p < 0.05; ** p < 0.01; **** p < 0.0001.

**Figure S4.**
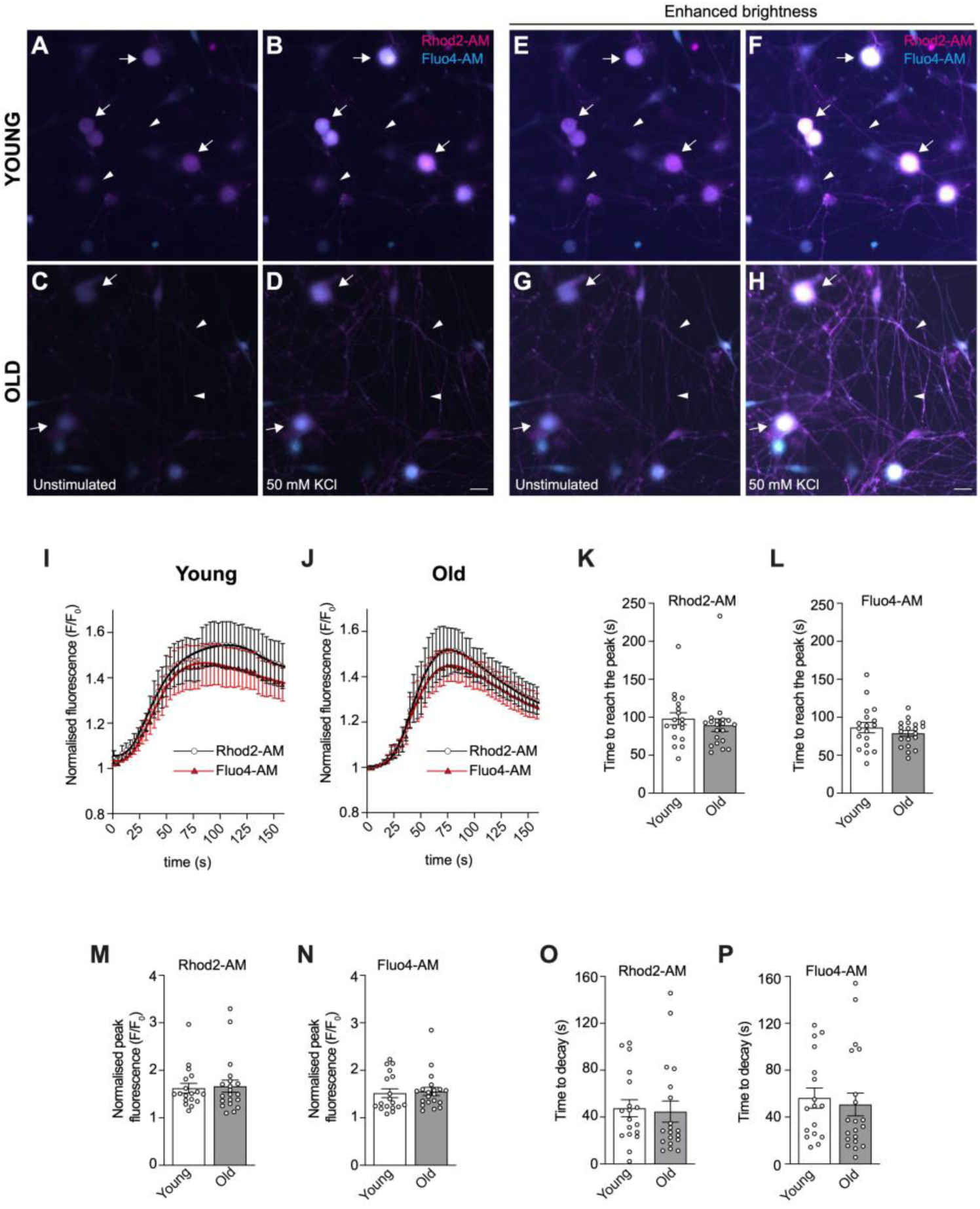
Mitochondrial and cytosolic calcium uptake are not significantly affected by age in cultured DRG neurons. **(A-H)** Representative images of DRGs cultured before (A,C,E,G) and after (B,D,F,H) stimulation with 50 mM KCl. Cells are co-stained with Rhod2-AM (magenta) and Fluo4-AM (cyan) for measuring the fluorescent signal of mitochondrial and cytosolic Ca^2+^, respectively. Neuronal depolarisation by KCl increases calcium uptake in cell bodies (arrows) and axons (arrowheads). Panels A-D are shown with enhanced brightness in E-H to highlight the response after treatment. Scale bars: 20 μm. **(I-J)** Traces indicate the average Rhod2-AM (black, circles) and Fluo4-AM (red, triangles) fluorescence intensity values in 18 young (I) and 20 old (J) neurons at individual time points (circles and triangles, respectively) normalised to the average fluorescence value before KCl stimulation (time 0). **(K-L)** Time to reach the peak, **(M-N)** normalised response peak and **(O,P)** decay time for cytoplasmic (Fluo4-AM) and mitochondrial (Rhod2-AM) calcium. Mann-Whitney tests (K,M-P) and an unpaired student’s t-test with Welch’s correction (L) showed no difference between conditions. DRGs were obtained from 3 young (P58-93, 3 males) and 4 old (P560-676, 1 female, 3 males) mice.

**Figure S5.**
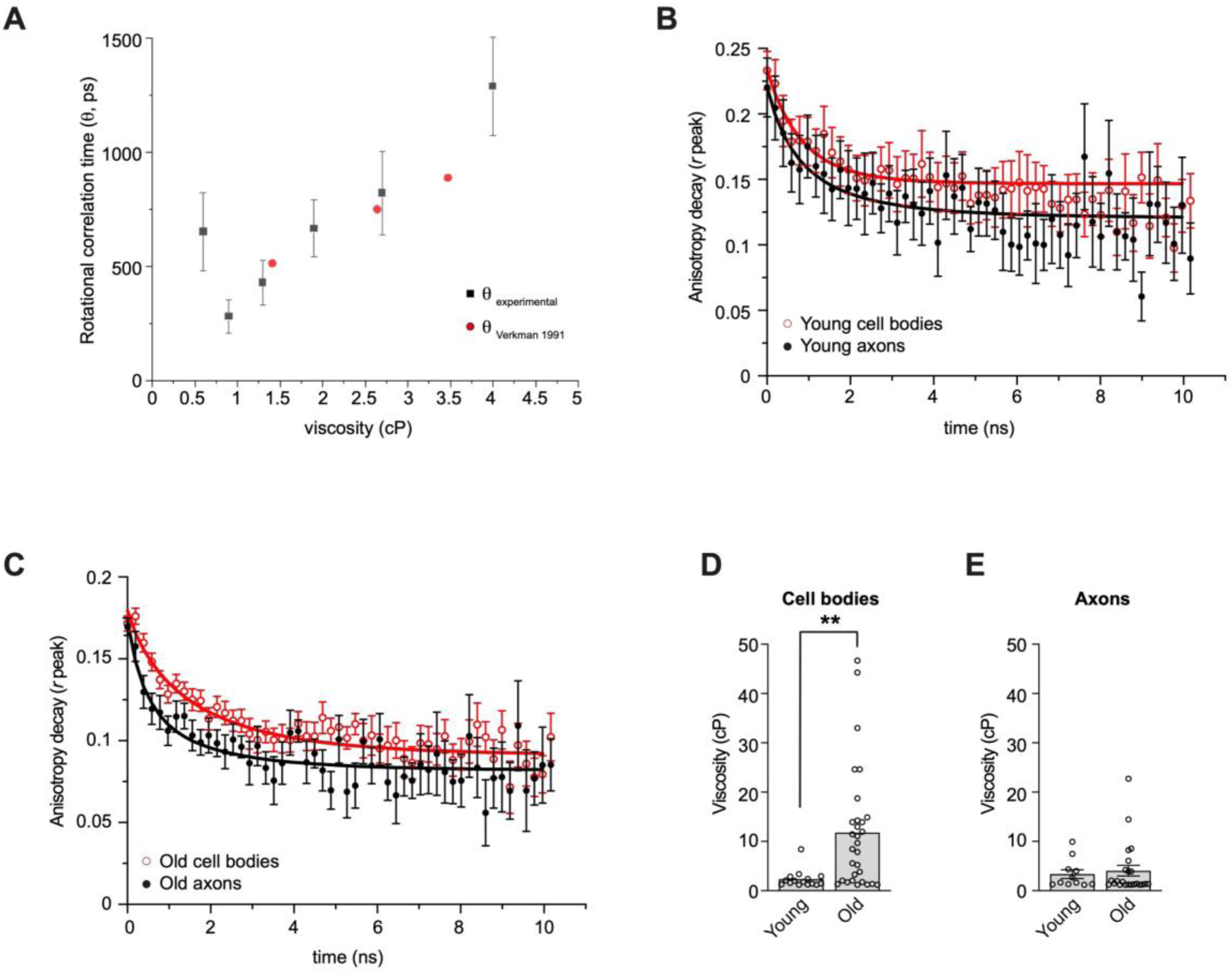
**(A)** Calibration curve of BCECF rotational correlation time (θ) calculated in accordance with the Stokes-Einstein-Debye equation. The experimental data were compared to historical data for BCECF θ as a function of viscosity (Verkman et al., 1991) and showed good agreement. **(B-C)** Fluorescence anisotropy decay and fit (see Methods, equation 3) of BCECF fluorescence obtained from perpendicular and parallel decays (according to equation 1, see Methods) from cell bodies and axons of young (B) and old (C) DRG neurons. Red and black lines depict exponential decay fit for cell bodies and axons, respectively. The anisotropy decay of young cells (B) is not significantly different between compartments, leading to similar viscosity values (Fig. 4D). The slower decay of cell bodies compared to axons in old DRGs is indicative of higher viscosity, as shown in Fig. 4E. Circles, individual time points. **(D-E)** Cytoplasmic viscosity increases during age in the cell bodies (D) but not axons (E) of DRG neurons. Circles depict cell bodies and axons imaged. Data are shown as mean ± SEM. Mann-Whitney test (D-E). ** p < 0.01.

**Figure S6.**
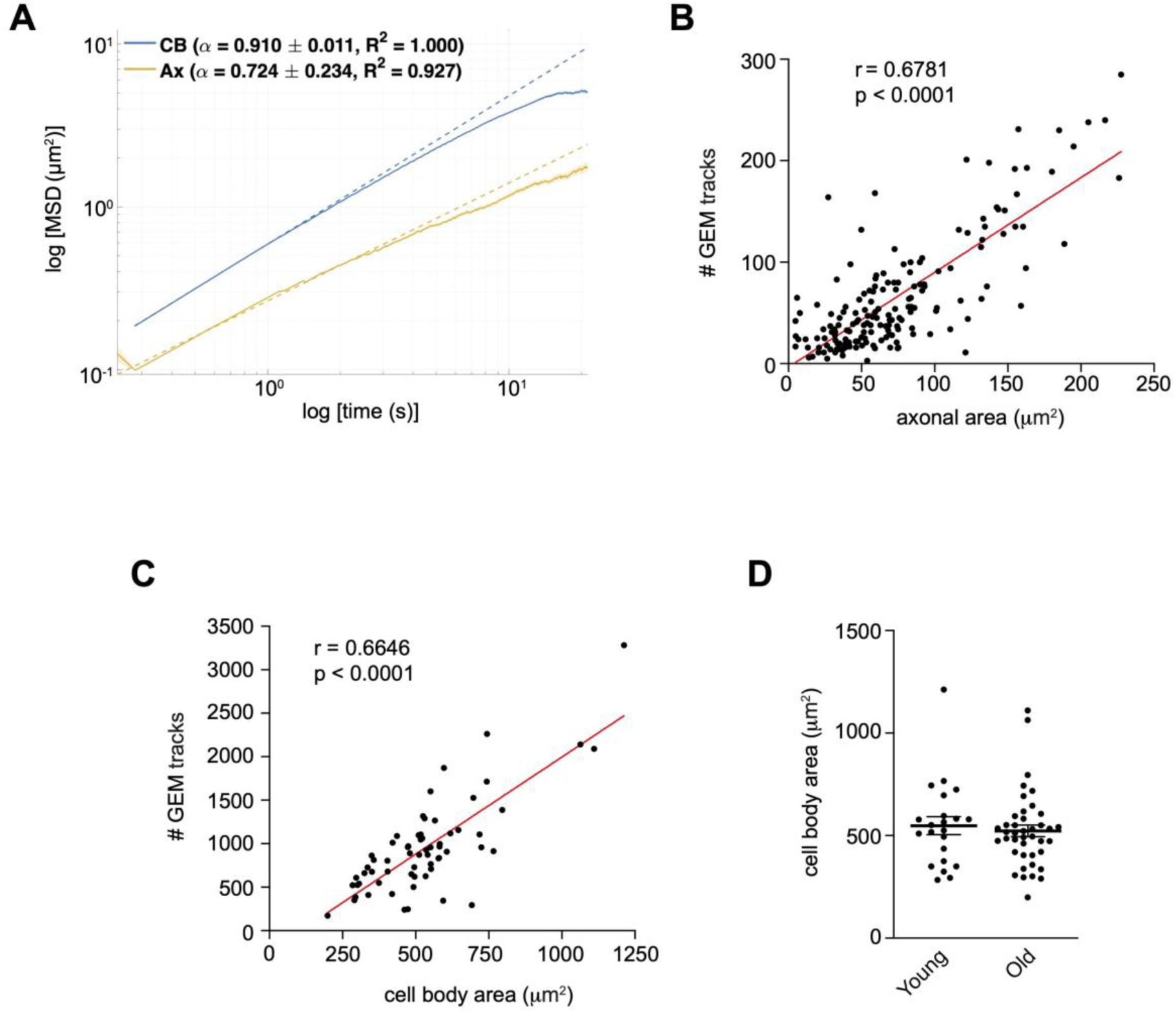
**(A)** Combined MSD of GEM tracks in cell bodies (CB, blues) and axons (Ax, orange). Data are plotted as mean ± SEM in a log-log format. Dashed lines are curve fit with anomalous exponent of 0.910 ± 0.011 (CB) and 0.724 ± 0.234 (Ax), respectively. **(B-C)** Positive correlation between total number of GEMs tracks analysed and axonal (B) and cell body (C) areas, suggesting that GEMs equally cover the full extent of the neuronal cytoplasm, independently of cell size. Circles represent individual axons (B) and cell bodies (C). Red lines, linear regression fit. Spearman rho test. **(D)** There is no significant difference between the mean area of the young and old cell bodies, indicating that cell size is unlikely to affect viscosity and diffusiveness. Circles represent individual cell bodies. Data are shown as mean ± SEM. Mann-Whitney test (p = 0.5148).

**Figure S7.**
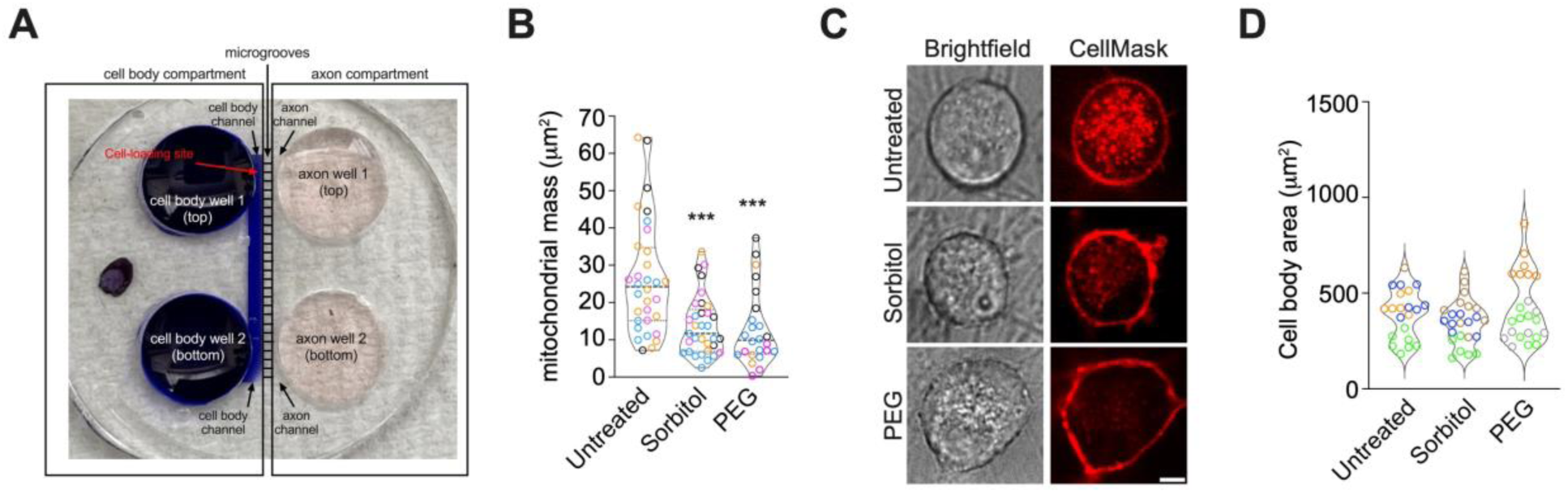
**A)** Microfluidic chamber (MFC) showing reliable fluidic separation between the cell body and axonal compartments. A blue dye was applied to the somal compartment at a lower volume than the connecting axonal compartment and incubated at 37°C for up to 5 h. Visual inspection confirmed that there is no dye leakage against the volume gradient. The same outcome was achieved after staining the axonal compartment. **(B)** Quantification of mitochondrial mass (see Methods) shows strong decline in sorbitol and PEG-treated DRGs. Datapoints in the violin plots represent individual microgrooves with each colour indicating a different animal (4 females, P45-86), Kruskal-Wallis with Dunn’s multiple comparisons test. **(C)** Representative images of DRG cell bodies that have been left untreated (top panels) or treated with sorbitol (middle panels) or PEG (bottom panels). The outline of the cell body defined by CellMask staining was used as a reference for the quantification of cell body area in (D). Scale bar: 5 μm. **(D)** Quantification of the DRG cell body area (see Methods) shows no significant change across conditions, unpaired t-tests with Welch’s correction (untreated vs sorbitol) and Mann-Whitney test (untreated vs PEG). Datapoints in the violin plots represent cell bodies from different MFCs, with each colour indicating a different animal (3-4 females, P45-86). *** p < 0.001.

**Table S1.**
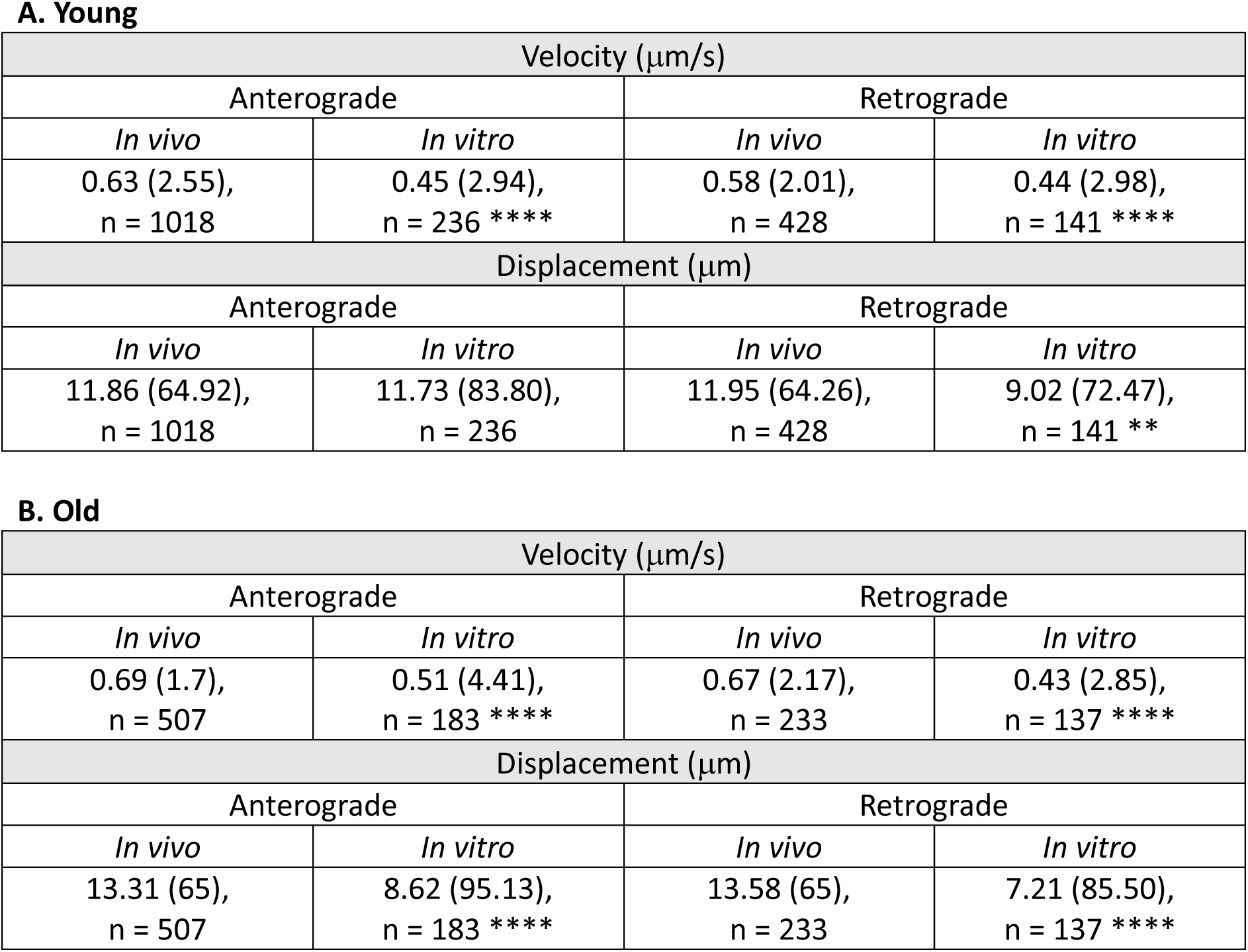
Median mitochondrial velocities and displacements are significantly higher in the sciatic nerve axons (*in vivo*) compared to DRG axons (*in vitro*) in all conditions except for anterograde displacement in young samples (A) where no difference was detected. Values are medians with, in brackets, maximum speeds and maximum displacements. n = number of mitochondria, Mann-Whitney test. ** p < 0.01; **** p < 0.0001. The data suggest that, while mitochondria are able to reach faster maximum velocities *in vitro*, on average they move more slowly *in vitro* than *in vivo*. Note that the displacement is calculated over 65 μm and 100 μm of axonal tract in the sciatic nerve and cultured DRGs, respectively, suggesting the calculated values may underestimate the significance between *in vivo* and *in vitro* displacements.

A. Young
| Velocity (μm/s) |  |  |  |
| --- | --- | --- | --- |
| Anterograde |  | Retrograde |  |
| *In vivo* | *In vitro* | *In vivo* | *In vitro* |
| 0.63 (2.55),<br>n = 1018 | 0.45 (2.94),<br>n = 236 **** | 0.58 (2.01),<br>n = 428 | 0.44 (2.98),<br>n = 141 **** |
| Displacement (μm) |  |  |  |
| Anterograde |  | Retrograde |  |
| *In vivo* | *In vitro* | *In vivo* | *In vitro* |
| 11.86 (64.92),<br>n = 1018 | 11.73 (83.80),<br>n = 236 | 11.95 (64.26),<br>n = 428 | 9.02 (72.47),<br>n = 141 ** |

Table S1
| Velocity (μm/s) |  |  |  |
| --- | --- | --- | --- |
| Anterograde |  | Retrograde |  |
| *In vivo* | *In vitro* | *In vivo* | *In vitro* |
| 0.69 (1.7),<br>n = 507 | 0.51 (4.41),<br>n = 183 **** | 0.67 (2.17),<br>n = 233 | 0.43 (2.85),<br>n = 137 **** |
| Displacement (μm) |  |  |  |
| Anterograde |  | Retrograde |  |
| *In vivo* | *In vitro* | *In vivo* | *In vitro* |
| 13.31 (65),<br>n = 507 | 8.62 (95.13),<br>n = 183 **** | 13.58 (65),<br>n = 233 | 7.21 (85.50),<br>n = 137 **** |

**Movie 1**

Representative time-lapse movie of mitochondria in axons of DRG neurons obtained from young (P92) and old (P676) mice and stained with MitoTracker Deep Red. Width: 73 μm. Movie plays at 20 frames per seconds and represents 2 min of real-time. Scale bar: 5 μm.

**Movie 2**

Representative time-lapse movie of mitochondria from young (P72) and old (P624) MitoMouse sciatic nerve axons. Width: 64 μm. Movie plays at 20 frames per seconds and represents 4:26 min of real-time. The ‘Smooth’ filter in Fiji/ImageJ was applied for presentation purposes. Scale bar: 5 μm.

**Movie 3**

Representative time-lapse movie of GEM particles in the cell body of a cultured DRG neuron. Movie plays at 20 frames per seconds and represents 30 seconds of real-time.

**Movie 4**

Representative time-lapse movie of GEM particles in the axon of a cultured DRG neuron. Width: 55 μm. Movie plays at 20 frames per seconds and represents 30 seconds of real-time. The ‘Smooth’ filter in Fiji/ImageJ was applied for presentation purposes.

**Movie 5**

Representative time-lapse movie of mitochondrial transport in MFCs acquired from axons of neurons in which the somal compartment was either left untreated (top movie) or was treated with sorbitol (middle movie) or PEG (bottom movie). Width: 80 μm. Movie plays at 20 frames per seconds and represents 1 min of real-time. The ‘Smooth’ filter in Fiji/ImageJ was applied for presentation purposes.

**Movie 6**

Representative time-lapse movie of mitochondrial dynamics in MFCs acquired from neuronal cell bodies. Top movie: untreated control; middle movie: cell body treated with sorbitol; bottom movie: cell body treated with PEG. Movie plays at 20 frames per seconds and represents 1 min of real-time. In the cell bodies treated with crowding agents, the mitochondria appear more clustered and less dynamic.

## AUTHOR CONTRIBUTIONS

JNS, FM, SR, EA, KN, IES, AV performed experiments, collected and analysed data; II, ILD, PPJ, SA analysed and interpreted data; ERR, JCM critically supported the experimental work; GY generated and provided reagents; JNS, KS, GS, AV supervised the work; JNS, KS, GS, AV were responsible for funding acquisition. AV wrote the manuscript with critical inputs from JNS, KS, GS. AV conceived and led the study. All the authors read and approved the manuscript.

## DATA AVAILABILITY STATEMENT

All relevant data that support the findings are within the paper and its supporting information files. All raw data are available from the corresponding author upon request.

